# Theory of branching morphogenesis by local interactions and global guidance

**DOI:** 10.1101/2021.08.30.458198

**Authors:** Mehmet Can Uçar, Dmitrii Kamenev, Kazunori Sunadome, Dominik Fachet, François Lallemend, Igor Adameyko, Saida Hadjab, Edouard Hannezo

**Affiliations:** Institute of Science and Technology Austria, Am Campus 1, 3400 Klosterneuburg, Austria; Department of Neuroscience, Karolinska Institutet, 17177, Stockholm, Sweden; Department of Physiology and Pharmacology, Karolinska Institutet, 17177, Stockholm, Sweden; IRI Life Sciences, Humboldt-Universität zu Berlin, Berlin, 10115, Germany; Ming-Wai Lau Centre for Reparative Medicine, Stockholm node, Karolinska Institutet, Stockholm, Sweden; Department of Neuroimmunology, Center for Brain Research, Medical University Vienna, 1090, Vienna, Austria

## Abstract

Branching morphogenesis governs the formation of many organs such as lung, kidney, and the neurovascular system. Many studies have explored system-specific molecular and cellular regulatory mechanisms, as well as self-organizing rules underlying branching morphogenesis. However, in addition to local cues, branched tissue growth can also be influenced by global guidance. Here, we develop a theoretical framework for a stochastic self-organized branching process in the presence of external cues. Combining analytical theory with numerical simulations, we predict differential signatures of global vs. local regulatory mechanisms on the branching pattern, such as angle distributions, domain size, and space-filling efficiency. We find that branch alignment follows a generic scaling law determined by the strength of global guidance, while local interactions influence the tissue density but not its overall territory. Finally, using zebrafish innervation as a model system, we test these key features of the model experimentally. Our work thus provides quantitative predictions to disentangle the role of different types of cues in shaping branched structures across scales.

## INTRODUCTION

Branching morphogenesis is a ubiquitous developmental process, where a number of morphogenetic events cooperate to give rise to complex tree-like morphologies. Branched structures are observed both at the level of multi-cellular organs, such as lung, kidney, mammary gland or vascular system [1–5], and at the level of single cells such as neurons [6] or tracheal cells [7]. A number of studies in the past decades have been devoted to understanding their design principles, with a particular emphasis on how given branched topologies and geometries can optimize properties such as transport and robustness [8–17].

A complementary question has been to understand the dynamical mechanisms through which branching complexity can arise during development. It has been shown in particular that branching morphogenesis proceeds via tip-driven growth and/or side branching events, which are shaped by combinations of deterministic and stochastic rules [4, 18–20]. Indeed, different cellular strategies have been demonstrated to regulate the final branching pattern, from stereotypic transcription factor expression [21], stochastic local rules [22, 23], mechanical forces and local reaction-diffusion mechanisms [2, 3, 24] to epigenetic mechanisms [25] and codes of cell adhesion molecules [19, 26]. In addition to these intrinsic mechanisms, branching morphogenesis is also controlled externally by a number of guidance cues from the environment [27–30], including chemical gradients (chemotaxis from diffusible factors or haptotaxis from substrate-bound adhesion or guidance molecules) or gradients in the mechanical stiffness of the environment [31, 32]. However, a theoretical framework to quantitatively assess the contribution of each intrinsic and/or extrinsic cue in shape, orientation and size of branched structures, as well as the relative roles of deterministic vs. stochastic factors during branching morphogenesis remains to be established.

Here, we combine numerical simulations with analytical theory to derive a comprehensive description of branching morphogenesis in the presence of internal self-organizing cues (such as self-avoidance of branches, stochastic exploration of space, and tip termination) and external guidance cues. Furthermore, we identify several metrics, including branch directionality, shape or efficiency of space filling, which are differentially affected by different model parameters. These metrics thus provide generic criteria, measurable from static data on the final branched structure, to distinguish different dynamical mechanisms at play during morphogenesis. Finally, we experimentally test our model in peripheral sensory system focusing on the branching of individual Rohon-Beard sensory neurons in the zebrafish caudal fin. Thus, we present a model where the combination of two simple parameters, for local self-interactions and global guidance, can synergize to generate complex branched structures both in two and three-dimensions.

## RESULTS

To analyze the influence of both the local selforganizing (intrinsic) cues and the global (extrinsic) guidance on the formation of branched structures, we first turned to a modelling approach inspired by the physics of branching random walks, which represents tips as particles undergoing both stochastic and deterministic elongation movements (which generates branches at speed *υ*), as well as stochastic branching events into two tips with probability *p_b_*. This type of model [20, 22, 33–35] has the advantage of coarsening many microscopic features of branching regulation (for instance that have been addressed via reaction-diffusion models [36, 37]) into simple sets of rules. In this work, we include both the possibility for global guidance via gradients quantified by a guidance strength *f_c_* (which acts as a deterministic force on tip motion) as well as local self-avoidance of neighbouring branch segments. Such self-avoidance can typically occur in neurons by cycles of contact-retraction when a tip touches a neighbouring branch of the same cell [20, 38], or in branched multicellular organs via diffusible molecules [39]. Here we concentrate on the morphogenesis of single neurons, and therefore model self-avoidance effectively by tips moving deterministically away from neighbouring branches of the same tree at strength *f_s_*. If the tip fails to reorient in close proximity to a neighbouring branch, it terminates its active growth and becomes irreversibly inactive (which we call termination/annihilation, see Fig. 1A-B for a schematic of the model).

**FIG. 1.**
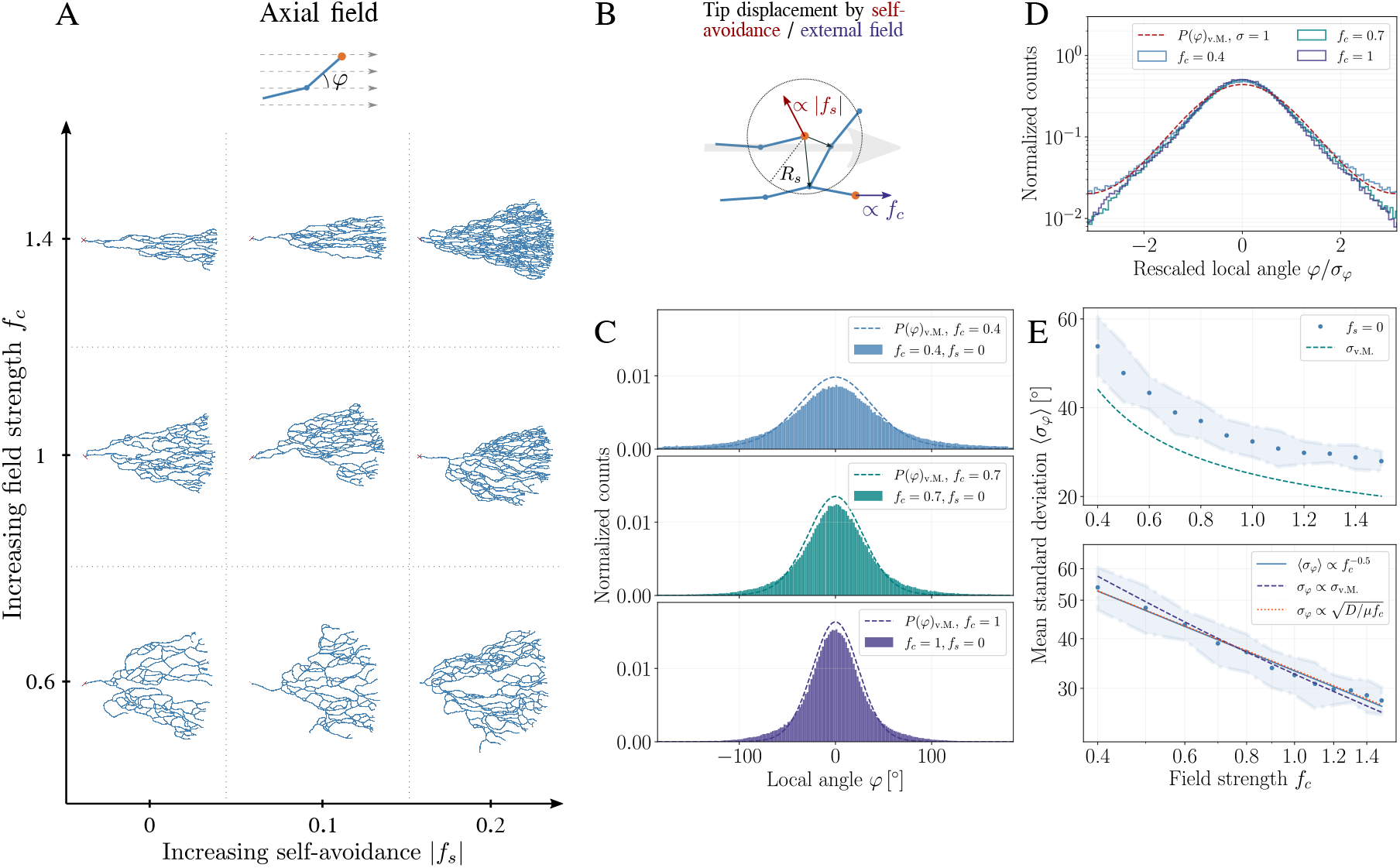
Morphology and alignment of branching structures in the presence of global guidance cues and local self-repulsion. (A-B) Schematic of the model and resulting branching morphologies. Top panels: we consider an active tip (orange node) which undergoes stochastic branching and elongation according to a local angle *φ* to make branch segments (blue solid lines), guided by an external field (linear guidance, dashed arrows). Self-avoidance or external field are implemented in the simulation by additional displacements of active tips of the branching network respectively by “sensing” neighboring branch segments (blue nodes) within a radius of repulsion *R_s_* (red arrow in B), or by a bias towards the external field (large gray arrow in B), respectively. The strength of local self-avoidance and external guidance are respectively determined by a factor |*f_s_*| or *f_c_*. Bottom panel: Morphology diagram of branching and annihilating random walks (BARWs) with linear (axial in one-dimension) external guidance obtained from simulations. Representative networks are displayed for different values of the external field strength *f_c_* and self-avoidance |*f_s_*|. (C) Probability distribution of tips growing with an angle *φ* for different values of *f_c_* and without self-repulsion (*f_s_* = 0) in the simulations (solid bars). These are well-approximated by the analytical predictions (dashed lines) following a von Mises distribution centered around zero and with single parameter 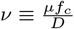. With increasing field strength *f_c_* the distributions become sharper, indicating better alignment of the branch segments with the external field. (D) Histograms of the local angle displayed in (C) rescaled by their corresponding standard deviations (SDs), showing that they all collapse onto the von Mises distribution with unit SD (dashed line), as predicted analytically. (E) Fluctuations in local angle *σ_φ_* decrease monotonically with increasing field strength *f_c_* (top panel), and exhibit a power-law relation (bottom panel) close to the scaling law predicted by the analytical theory (dashed line) and to the scaling law 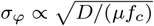) (dotted line).

To determine how the morphology and shape of a branching structure is affected by local intrinsic vs. global extrinsic cues, we concentrated on the two key parameters of local self-avoidance *f_s_* and of global gradient *f_c_*, and asked whether each gave rise to qualitatively different types of morphologies. Indeed, building a phase diagram of branching morphologies revealed key differences: in the presence of an external, axially oriented (linear) gradient, branched structures adopt triangular shapes, branching in a cone-shape with a well-defined angle that becomes smaller for increasing guidance strength *f_c_* (Fig. 1A, bottom panel). On the other hand, changing the self-avoidance strength *f_s_* gave rise to denser, as well as visually more aligned branches, but did not markedly change the overall shape.

To back the qualitative insights of this phase diagram more quantitatively, we sought to develop an analytical theory of branching via external guidance, which falls under the class of branching and interacting random walks [22] with external bias (external field). Starting from a microscopic description of branching and elongation events, we derived a (continuum) Fokker-Planck equation for the tip growth and branching under the influence of external field guiding elongation (as described in detail in SI Note). In particular, we obtained an equation for the time evolution of the probability of a tip to grow in a given direction, denoted by an angle *ψ* relative to the polarity of the external field. This direction is subjected to two types of random fluctuations: a gradual one arising from the randomness of tip elongation and abrupt changes arising from branching events (see section S1 of the SI Note for details on the methods to treat these two types of fluctuations based on Ref.[40]). These two sources of stochasticity are integrated into a diffusion coefficient *D* while the external guidance gradient acts as an effective velocity term *μf_c_* in the Fokker-Planck equation:

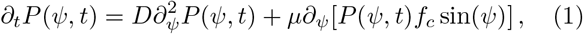

which reflects a sinusoidal reorientation of the active tips by the external field [41]. Importantly, this equation attains a steady-state solution (*∂_t_P*^st^(*ψ, t*) = 0) that is largely independent of the form of the external field. This solution predicts that the alignment of angles with respect to the polarity of the external field will be determined by the von Mises distribution (circular normal distribution [42]):

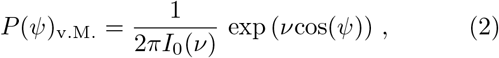

with a concentration parameter given by *ν* ≡ *μf_c_/D*, and *I*_0_(*ν*) is the modified Bessel function of the first kind of order zero. The fluctuations in the angular alignment as determined by the variance will thus follow a universal scaling approximately given by *σ*^2^ ∝ *D/μf_c_* that underlines the relative contribution of the local noise to the external guidance. For an axial (linear) potential parallel to the horizontal axis, for instance, the above solution applies to the distribution of the local angles *φ* of the branch segments (Fig. 1C). In a radial external field, however, the alignment of a branch is determined by the angle difference *ψ* ≡ *φ* – *θ* between its local angle and its angle *θ* with respect to the origin of the external field, and thus *ψ*, rather than *φ*, is predicted to follow the von Mises distribution (see Fig. S1A for the alignment angles in different external fields). Comparing these analytical criteria with the numerical simulations led to excellent agreement without using any fit parameter (Fig. 1C-E).

To test these predictions, we examined the morphology of sensory neurons during zebrafish fin innervation as a model system, as it has several advantages: i) it is a simple quasi two-dimensional (see Movie S1) and transparent system, facilitating imaging and reconstruction, ii) the innervation pattern is complex, with tens to hundreds of branches per neuron, and iii) multiple axons arise from dorsal part of spinal cord and start branching out in a simple geometry, *i.e*. a roughly semi-circular region (Fig. 2A). To segment and reconstruct single branched neurons, we used sparse labelling of mCherry positive neurons (see Fig. S7) at 5 days post-fertilization (a time when neurons are functional and the fish is able to swim), and skeletonized the manually traced filaments to generate hierarchical tree topologies (see section S3 of the SI Note for details). Interestingly, we found that these neurons, although all appearing to grow radially towards the outer edge of the fin, were highly stochastic and heterogeneous both in shape, size, and morphology (Fig. 2C, Fig. S8), and covering domains of highly different sizes. This hints at a highly stochastic pattern of fin innervation, as expected in our branching random walk model when we adapted it to a radial external field (Fig. 2B). Qualitative comparisons with different stochastic simulations with identical model parameters revealed good visual agreement (Fig. 2C-E), exhibiting similar stochasticity in shape, angles, topology and size of neuronal trees, as seen in the experimental data. Furthermore, few crossovers between branches could be observed with terminal tips residing all over the neuronal structure close to neighbouring branches (Fig. 2A,C), as qualitatively expected in the framework of branching and annihilating random walks. Finally, and more quantitatively, we extracted the branch length distribution across neurons, and found that it was very wide (with branches of all lengths seen in data) and well-described by a simple exponential, as predicted by a stochastic branching process (Fig. 2F). Altogether these key features supported the applicability of our theory of branching and annihilating random walks to the experimental dataset.

**FIG. 2.**
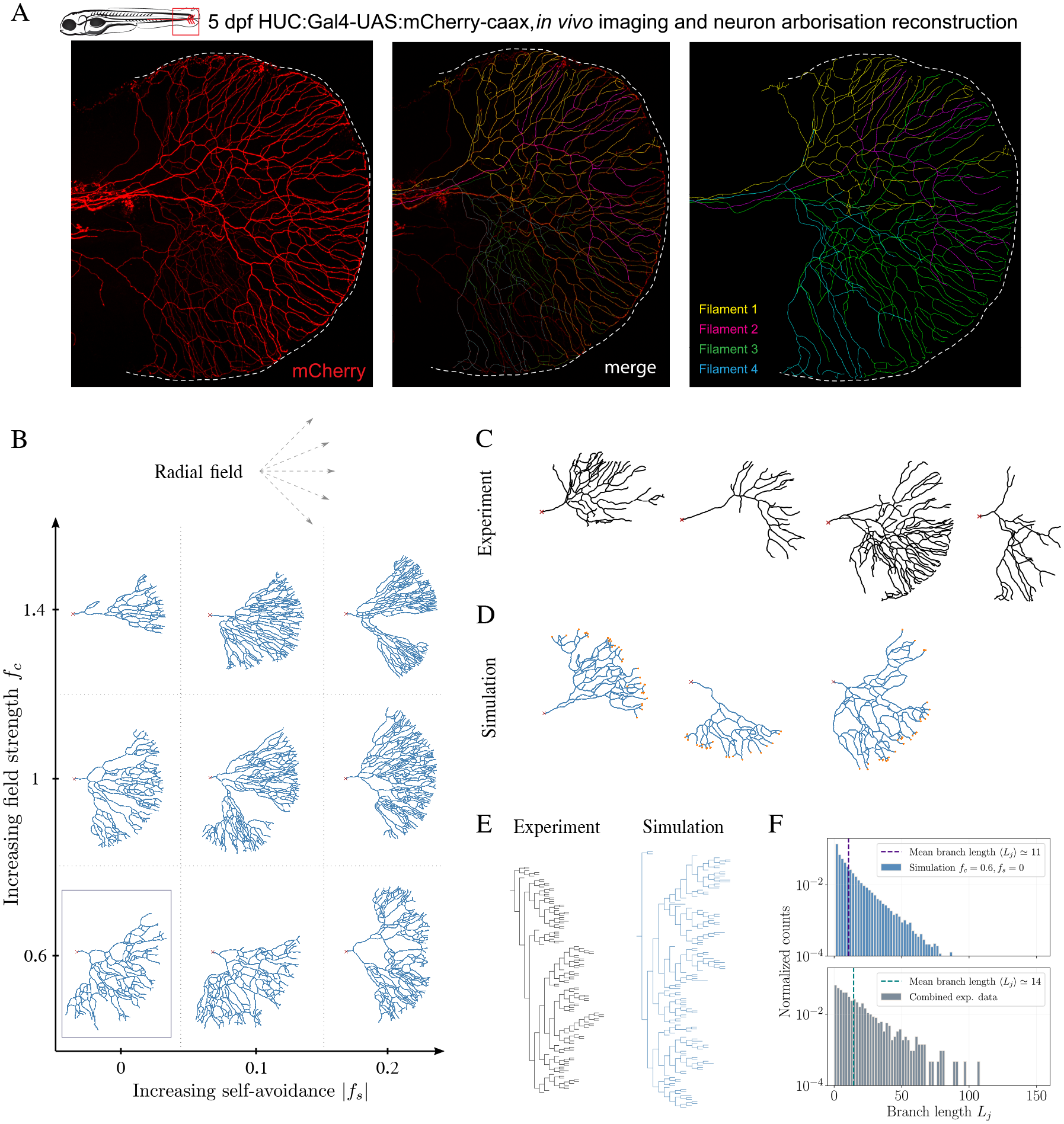
Branching and annihilating random walks (BARWs) with radial guidance cue reproduces qualitative features of zebrafish caudal fin innervation. (A) Development of the zebrafish nervous system and innervation of the caudal fin (boxed area in the top left cartoon) 5 days postfertilization. (Left) Confocal image of neuronal cell membranes in the caudal fin imaged via red mCherry fluorescence (HUC:Gal4-UAS:mCherry-caax). Imaged Rohon-Beard sensory neurons exhibit a clear directionality towards the fin edge (indicated by the dashed white lines). (Middle) mCherry labelled neurons (red) visualized together with the manually reconstructed filament trees (different colors overlaid), showing the overall faithfulness of the reconstructions. (Right) Different neuronal trees color-coded for visualization. (B) Morphology of branched structures with the same model as in Fig. 1, but in a radial external potential (dashed arrows, top) obtained from simulations for different values of the external field strength *f_c_* and self-avoidance |*f_s_*|. (C-D) Simulations with an intermediate external field strength (*f_c_* = 0.6) and no self-avoidance (*f_s_* = 0), corresponding to the boxed region in the morphology diagram (B), capture the overall directionality observed in reconstructed networks (four representative neurons, red cross indicating “origin” of the axon, C), but also show some stochasticity in the final network structure as in the data. Active tips of the simulated branching networks are highlighted in orange. (E) Comparison of tree topologies between two exemplary networks obtained from simulations (right) and experiments (left), emphasizing the common stochasticity and heterogeneity in subtree sizes. (F) Branch lengths *L_j_* (in normalized units) obtained from simulations (top) and experiments (bottom) are distributed exponentially, as predicted by our theory of stochastic branching.

To go further, we turned to the quantitative structure of the branching patterns, and first analyzed the distribution of branch angles in the data. As predicted by Eq. (2), we expected the distribution of the angle difference *ψ* to decay with a variance scaling as *D/μf_c_* (see Fig. 3B-D for the distributions *P*(*ψ*) obtained from simulations and analytical theory). Comparing theory and experimental data revealed very good agreement, with both single neuron distributions (see Fig. S10 in SI Note for the in dividual distributions) and distributions averaged across all data (Fig. 3E) closely following the predicted scaling of a von Mises distribution. Importantly, the single free parameter in this fit (*i.e*. the variance of the distribution) allows us to estimate *μf_c_/D*, and thus the relative strength of the global/extrinsic guidance compared to local stochasticity (see section S3.6 in SI Note for details on the measurements of the other parameters, in particular the estimation of the branching probability and branch length). Interestingly, we find intermediate values of *D/μf_c_* ≃ 0.35, arguing that neuronal morphology is shaped by a combination of both factors.

**FIG. 3.**
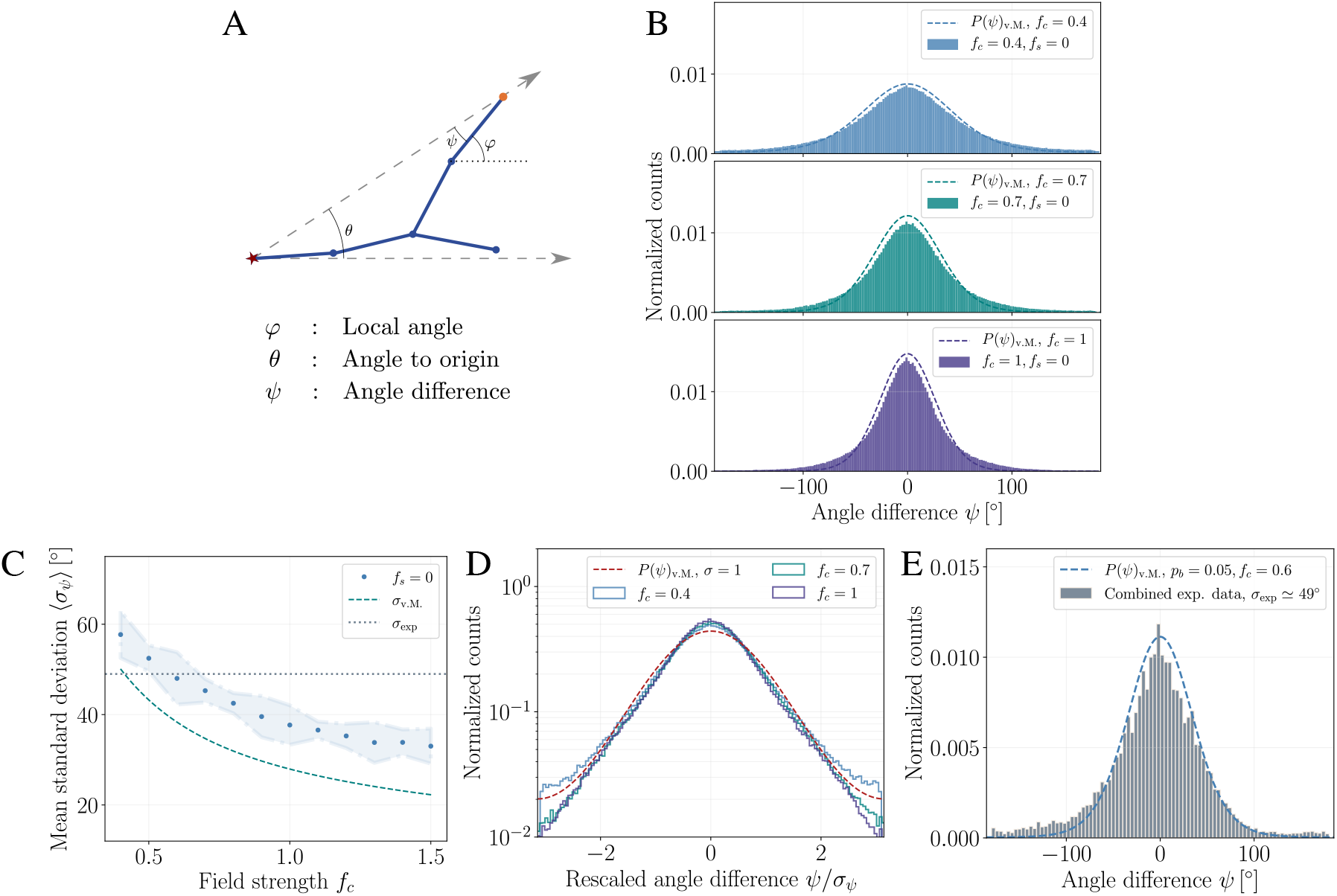
Continuum model predicts the alignment of branch segments along the external field both for simulation and experimental data. (A) Representative cartoon of branch segments in a radial external field (dashed arrows) highlighting three distinct angles: The *local angle φ* of a branch segment with an active tip (orange node), the *angle to origin θ* (denoted by the star symbol, which determines the extrinsic guidance direction at this point), and the *angle difference ψ* ≡ *φ* – *θ* (which tends to be minimized by extrinsic/global guidance). (B) Normalized histograms of the angle difference *ψ* for different values of *f_c_* and without self-avoidance (*f_s_* = 0). The histograms (solid bars) are well-approximated by von Mises distributions (dashed lines) as for the local angle *φ* in a linear gradient (Fig. 1), and as predicted by the continuum model. (C) Mean standard deviations (SDs) 〈*σ_ψ_*〉 of the angle difference *ψ* obtained from simulations are close to the mean SD of the experimental data (dotted horizontal line) for an external field strength of *f_c_* = 0.6. For comparison, the scaling of SD as a function of *f_c_* predicted by the von Mises distribution is displayed (dashed line). (D) Histograms of angle difference *ψ* rescaled by their corresponding SDs *σ_ψ_* (solid lines) are well-approximated by the von Mises distribution with unit SD (dashed line). (E) Angle difference distribution obtained from the experimental data from *n* = 8 networks (solid bars) compared with the von Mises distribution predicted by the theory for an external field strength of *f_c_* = 0.6 (dashed line). The latter value is inferred from the matching of theoretical and experimental values of the SDs displayed in (C), and no other fit parameter is used.

Such extrinsic guidance provides a simple theoretical mechanism to restrict neuronal growth to a domain characterized by a well-defined opening angle 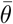. Turning to experiments, we found that reconstructed neurons were typically also characterized by such angle, which we estimated as 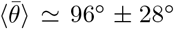. Theoretically, the average opening angles 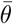 decreased monotonically with increasing field strength *f_c_* (see Fig. 4A for an illustration) with strikingly similar values both in the presence and absence of self-repulsion (see Fig. 4B). Using a simple geometric argument we could approximate these opening angles by:

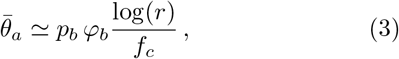

where *r* is the radial distance of the furthermost branch from the origin of the network (fixed by the maximal time of network growth). With this approximation, we could fit the numerical data by using a single fit parameter *φ_b_* (see section S2.5 in SI Note for further details). From the fitted value of the external field *f_c_* = 0.6, we predict an opening angle of 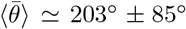 (mean ± SD). Although this overestimates the experimental value, we note that this prediction is based on a perfectly radial gradient in 360° without boundary, *i.e*., assuming neurons can branch backwards. When we confined the theory to a 180° hemispherical region, which seems to reflect the experimental geometries (Fig. S8), we obtained average opening angles of 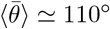, much closer to the data.

**FIG. 4.**
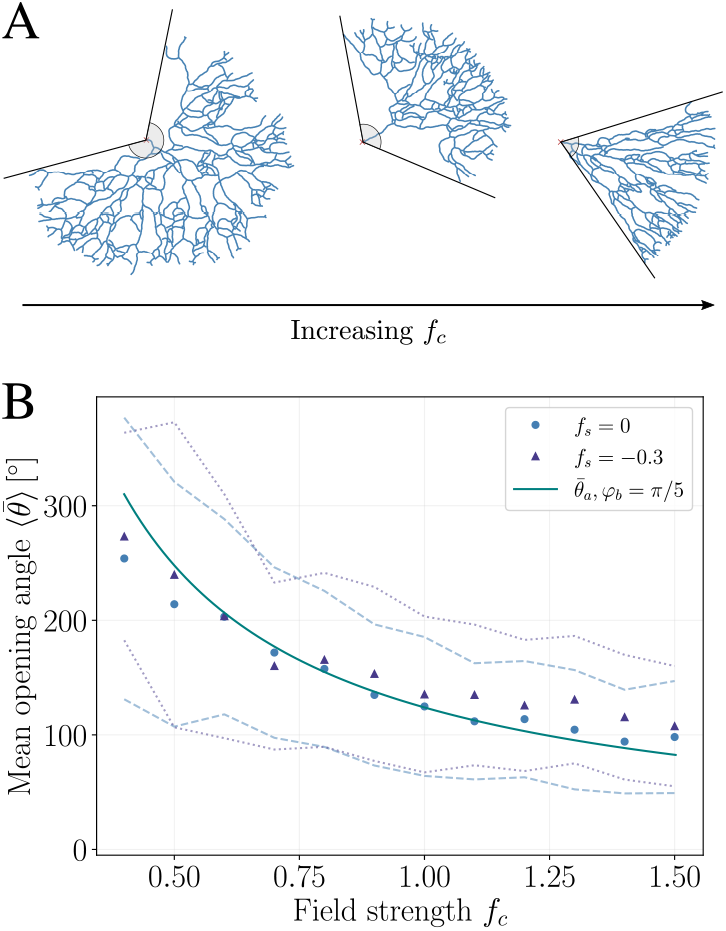
The strength of extrinsic guidance determines the territory of the branched structures in simulations, with minimal influence from local self-avoidance. (A) Representative simulation snapshots highlighting the changes in the opening angle 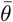 (a proxy for territory size in a radial geometry, gray circular segments) with increasing external field strength *f_c_*. (B) Mean opening angles 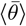 of branched networks decrease monotonically with increasing external field strength *f_c_*, and have similar values for networks with zero (*f_s_* = 0, circular markers) or with strong (*f_s_* = −0.3, triangular markers) self-avoidance. Errors of the averages are determined by the standard deviations and highlighted by the dashed and dotted lines for *f_s_* = 0 and *f_s_* = −0.3, respectively. The monotonic decrease of 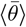 obtained from simulations is well-described by the analytical approximation 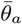 (solid line), see Eq. (3), with the fit parameter *φ_b_* = *π*/5.

Although the existence of an external gradient has not yet been characterized in the zebrafish fin, we note that other features from the comparison between experimental and theoretical data argue in favour of it. For instance, even though self-avoidance can lead locally to aligned branches, these branches grow isotropically in any direction without global cues (see section S2.3.1 and Fig. S3 in SI for a brief illustration). This is in particular true in low-density regions (which occur stochastically in the simulations), where fewer branches would lead to weaker repulsive cues, and consequently, in the absence of an external gradient, would result in tips deviating from the radial direction. However, examining the data revealed that this did not occur: even in sparse branching regions (*e.g*. Fig. 2B-D), branches appear as directional towards the fin periphery as in dense branching regions. Furthermore, sparse neurons also showed the same angle alignment distribution as dense neurons in the data (Fig. S10C), in contrast to what would happen in the absence of external guidance.

Finally, we sought a quantitative metric which could distinguish between networks with weak or strong self-repulsion *f_s_* after having estimated *f_c_*. Visually, our phase diagram of neuronal morphology confirmed the intuitive idea that larger self-avoidance *f_s_* should allow for denser networks. Furthermore, we tested the less intuitive effect of self-avoidance on the efficiency of space tiling [43, 44], by quantifying the fractal dimension *d_f_* of the branching networks (box-counting method, Fig. 5A) as a function of model parameters. We found that self-avoidance markedly improved the space-filling properties of the branching networks (see Fig. 5B, a fractal dimension close to *d_f_* = 1 is expected for very sparse structures, while a fractal dimension of *d_f_* = 2 corresponds to full tiling of space). Then, we again turned to the experimental data to ask whether these signatures could be observed. Because the branching rate/number showed variability across samples, we first explored this effect, and found a positive correlation between mean branch probability in a neuron and its fractal dimension (Fig. S11), as expected. Focusing on the four densest networks to remove this confounding effect, we found that measuring fractal density in experiments yielded robust power-laws as predicted by the simulations (Fig. 5C), with a typical exponent in the range of *d_f_* ≃ 1.55 ± 0.04 (mean±SD). This is consistent with our computational screen for relatively small values of self-avoidance (in the range of |*f_s_*| = 0 — 0.1), a feature which was confirmed by comparing absolute densities between model and data (Fig. 5D-F). This argues that although we cannot exclude a small contribution of self-repulsion in locally aligning branches, global external guidance cues play a dominant role in shaping these neuronal structures.

**FIG. 5.**
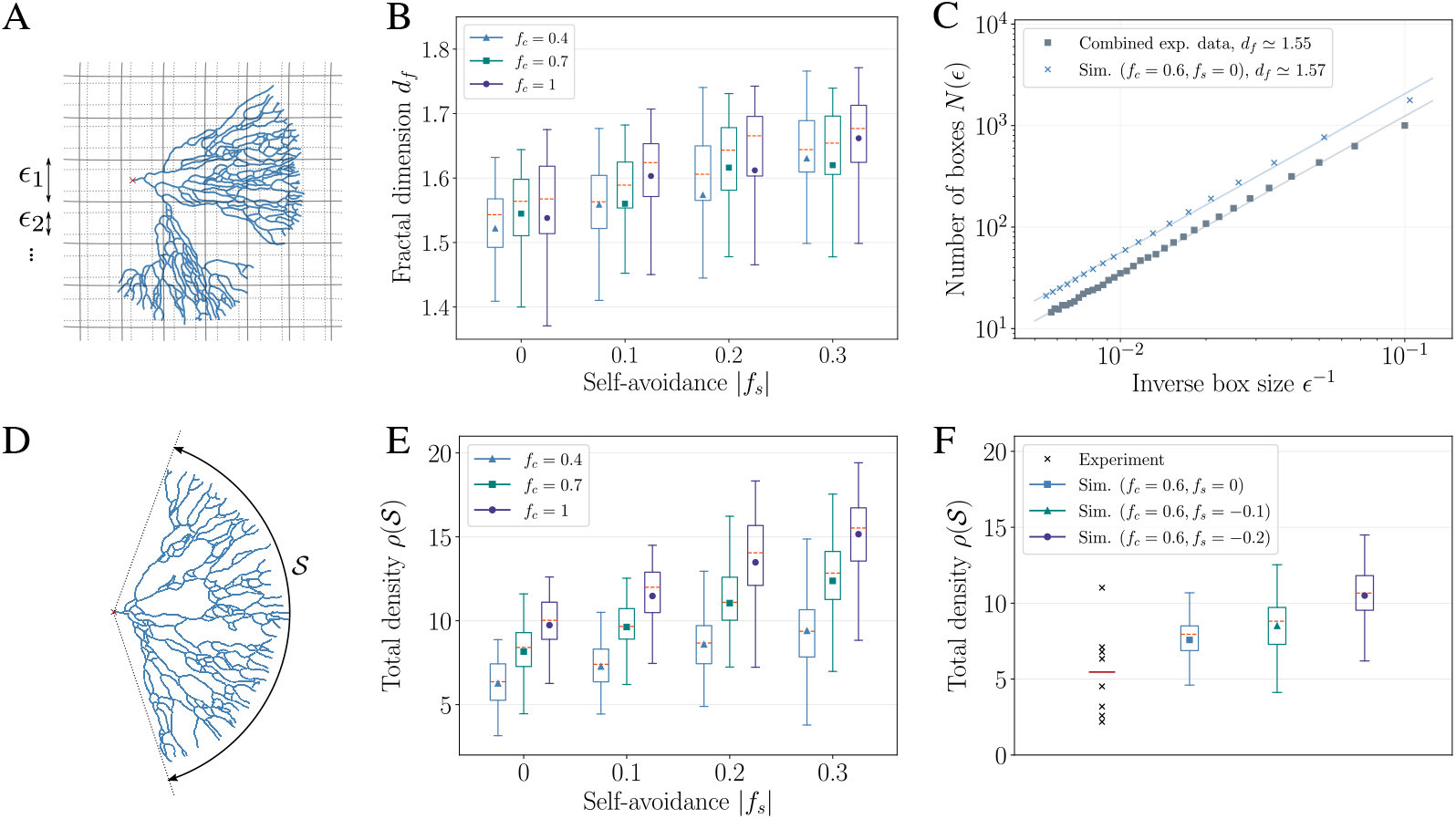
Effect of self-avoidance and external guidance on branching density and space-filling properties. (A) Fractal dimension of the networks estimated by the box-counting method: Boxes of decreasing sizes *ϵ* are used to count the total number of boxes that include at least one skeletonized node. (B) Mean fractal dimensions obtained from the box-counting method increases from 〈*d_f_*〉 ≃ 1.52 to 〈*d_f_*〉 ≃ 1.67 with increasing self-avoidance (*f_s_* = 0 to *f_s_* = −0.3), whereas large changes in the external field strength (from *f_c_* = 0.4 to *f_c_* = 1) have a smaller effect on the mean values. (C) Combined experimental data from the densest *n* = 4 networks (circular markers) showing a clear power-law signature as theoretically predicted. We find a fractal dimension of *d_f_* ≃ 1.55, close to the theoretical value *d_f_* ≃ 1.57 obtained from the combined data from simulations with *f_c_* = 0.6 and *f_s_* = 0 (crosses). In all box plots, mean and median values are given respectively by the plot markers and horizontal dashed lines (orange), and whiskers indicate 1.5 interquartile range. (D) Average density 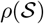 of a branched network (ratio of the number of branches to the arc length 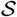 spanned by the network). (E) Densities 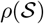 of the simulated networks increase markedly both for increasing external field strength *f_c_* and self-repulsion strength |*f_s_*|. (C) Densities 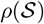 obtained from experimental data for *n* = 8 filaments (crosses) compared with densities obtained from simulations for *f_c_* = 0.6 and *f_s_* = 0 (blue box), *f_s_* = −0.1 (green box), and *f_s_* = −0.2 (purple box). Mean density of the experimental data (red horizontal line) is on the lower end of the densities obtained from simulations even for low repulsion, indicating a small value for the parameter *f_s_*.

## DISCUSSION

In this work, we have derived an analytical theory, backed by stochastic numerical simulations, of branching morphogenesis under both local cues-such as repulsion, branching, and termination as well as global guidance from external cues. Each of these factors can be tuned to create a variety of complex branched structures. To systematically classify these, and try to understand analytically how each parameter impacts the final structure, we derived a continuum Fokker-Planck theory, which enables us to coarse-grain the parameters of the numerical simulation (branching angles, branching rate, stochasticity in elongation, external guidance strength) into a few relevant coefficients at the macroscopic level. Through this, we have identified a number of generic features in the final branched structures. For instance, a combination of branching/elongation stochasticity in the presence of global guidance cues generically gives rise to well-defined scaling laws for the alignment of branch angles, which only depend on the geometry of the problem, with a variance that can be used to extract the relative contribution of each effect. Space-filling properties such as fractal dimension on the other hand are strongly optimized by local parameters such as self-repulsion.

Our approach here is based on a minimal model to understand the growth of branched structures from simple rules (elongation, branching, guidance, avoidance) within a statistical physics framework. At smaller scales, one would need to take a number of features into account, for instance the specifics of axonal/substrate mechanics during neuronal growth [31, 32], to understand what regulates mechanistically each of the parameters that we use in the model. A strength of our “mesoscopic” approach is that it extracts a small number of such coarsegrained parameters, to identify which ones are key at the scale of the overall branching pattern, and thus guiding subsequent, more detailed modelling. Our proposed framework builds upon previous simulations of stochastic branching morphogenesis, which had considered local cues such as branching and repulsion [20, 22, 33, 35]. We find that adding global extrinsic guidance-a key element in different contexts to break the isotropy in tissue growth-in the model gives rise to significantly different dynamics, enriching the phase diagram of possible branching patterns. Furthermore, in addition to the computational/numerical features of this framework, we provide a continuum theory for branching morphogenesis guided by extrinsic cues, which enables us to make simple but generic predictions on testable experimental metrics such as the orientation of branch segments.

To begin to test this theory, we have examined the innervation of the zebrafish fin, which proceeds in a simple quasi-2D radial geometry, and, despite the local stochasticity, displays a strong overall radially-oriented bias towards the fin edge. Quantitative reconstructions of several neurons allowed us to test a number of metrics predicted by the theory in the experimental data, such as the distributions of branch lengths and branching angles, or the space-filling properties of individual neurons. In particular, the observation that fin neurons exhibit a clear directional bias with rather well-defined angles can be readily explained in our framework by simply emerging from a global/extrinsic guidance cue which directs single neuronal tips towards the outer edge of the fin. Identifying such an interaction would be a natural next step. It has been shown for instance in the zebrafish pectoral fin that molecules such as BMP or Smoc1 are patterned in a graded way towards the edge during morphogenesis [45], and that innervation of the pectoral fin exhibits a strong variability in sensory neuron morphologies [46]. Overall, a global guidance cue would provide a minimal/complementary explanation to the more involved mechanism of repulsion/tiling between branches of different neurons [47]. Such hetero-avoidance would lead to more refined boundaries between neighboring neurons and could in particular play a role in reducing the domain angles occupied by the individual neurons.

This theoretical framework, although we have applied it here to a specific geometry in neuronal branching, is highly general and could be applied to any branching structure such as in angiogenesis or the branching of epithelial organs, where similar questions on external guidance vs. local self-organization rules arise [48–50]. Similarly, it has recently been proposed that these types of stochastic tip interaction mechanisms are conserved for the morphogenesis of various filamentous organisms such as plants or Fungi [51]. Understanding quantitatively the relative contribution of each mechanism is also of key importance for the morphogenesis of branched mammalian organs: Mammary gland or late-kidney morphogenesis have been proposed to follow a simpler form of these stochastic models in the absence of external guidance [22], although kidney morphogenesis has been suggested to require larger amount of self-avoidance strength (denoted by the parameter *f_s_* in our framework) at early stages to avoid premature termination [52]. This hints at a potentially broad applicability of our framework in a wide number of systems, which would be a next step for future research.

## Acknowledgements

We thank all members of our respective groups for helpful discussion on the manuscript. The authors are also grateful to Prof. Abdel El. Manira for support and sharing Tg(HUC:Gal4;UAS:Synaptohysin-GFP), to Haohao Wu for discussion, to the zebrafish Core facility and Biomedicum Imaging Core (BIC, with support from the Karolinska Institutet) for technical support. This work received funding from the ERC under the European Union’s Horizon 2020 research and innovation programme (grant agreement No. 851288 to E.H.) and under the Marie Sklodowska-Curie grant agreement No. 754411 (M.C.U.), and by Ming Wai Lau Foundation (F.L.); S.H. is supported by the Swedish Research Council.

## Author contributions

Project initiation: S.H.; Project supervision: S.H. andE.H.; Project conceptualization: S.H., E.H., M.C.U.; Experiments and reconstructions: D.K., K. S.; Theoretical model: M.C.U.; Data analysis: D.K., M.C.U., D.F.; Resources and methodology: F.L., I.A.; Writing, original draft: E.H., M.C.U.; Writing, review and editing: S.H., together with inputs from all authors.

## Supplementary Information

In this Supplementary Note, we provide additional details on the model derivation, on the numerical simulations of branching with repulsion and external guidance, as well as on the data collection, analysis and fitting to the model.

### S1 Derivation of the continuum model

In the continuum description, we concentrate on incorporating external guidance into a model of branching and annihilating random walks, in order to make simple predictions for the orientation of branches/tips during branching morphogenesis.

#### S1.1 Alignment angle as a random variable

The growth of a branching tissue with active tips in an external potential can be in principle modelled in the framework of a *biased and persistent* random walk in two dimensions. However, in such a framework, a full expression for the time evolution of the probability density *p*(**r**, *t*) for an active particle to be at location **r** at time *t* in general cannot be obtained in a closed form [1], and will involve several bias and reorientation terms due to different frequencies arising from elongation vs. branching events. We therefore sought to reformulate the problem in a simpler framework for a single random variable. After realizing that if we focus on the *alignment* of branch segments with an external field, rather than their exact position and polarity, we could restate the problem as finding how much the local angles of the segments diverge from the local direction of the external field. To avoid confusion, we note that by *branch segments* we refer to the elementary vectors of a fixed size *ℓ*, corresponding to the discrete steps taken by active tips, that are linked together consecutively to define the branches of a tree.

Angular alignment of a branch segment is quantified by different angles for the two cases of external fields discussed in the main text: (i) For a horizontally oriented, axial external gradient (directed towards the positive *x* direction), the *local angle φ* of a branch segment with respect to the horizontal axis determines its alignment along the field, (ii) For a radial external field emerging from a central point of origin (which is relevant for the experimental geometry of the zebrafish fin), however, the angular alignment will be given by the *angle difference ψ*:

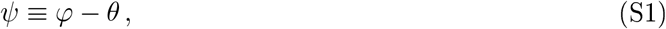

where *θ* denotes the angle of the active tip coordinates with respect to the origin of the external field. Fig. S1(A) illustrates the alignment angles for the two different external potentials. Note that, the axial potential is a limiting case of a radial field where the origin of the external field is located at *x* → −∞. This choice then corresponds to setting *θ* = 0 where the alignment angle *ψ* becomes equal to the local angle *φ*. Because of its generality, we will derive our model for the radial case using the angle difference *ψ* as the alignment angle in the following. Expressions for the axial case can then be obtained simply by setting *θ* = 0.

##### S1.1.1 Effect of an external field on the angle difference *ψ*

We now consider an external field that will influence the directionality of growing tips. At each step, the angle of a given active tip will be modified both stochastically via the randomness associated to elongation direction and branching, but also deterministically via guidance from the external field. For a branching network invading a circular region (in two-dimensions) we can define a radial potential that will displace the tips towards the point *P* with coordinates given by the vector **r** + *f_c_* **p**_*c*_, where *f_c_* determines the strength of the external field, **r** is the distance vector of length *r ℓ* between the active tip and the field origin, and **p**_*c*_ is a vector of length *ℓ* parallel to **r**. We note that we discretise the problem by having tip elongation by a small length *ℓ* = 1 (relative to the overall size of the network) without loss of generality. After such a displacement by the external field, the angle difference is given by

**Figure S1:**
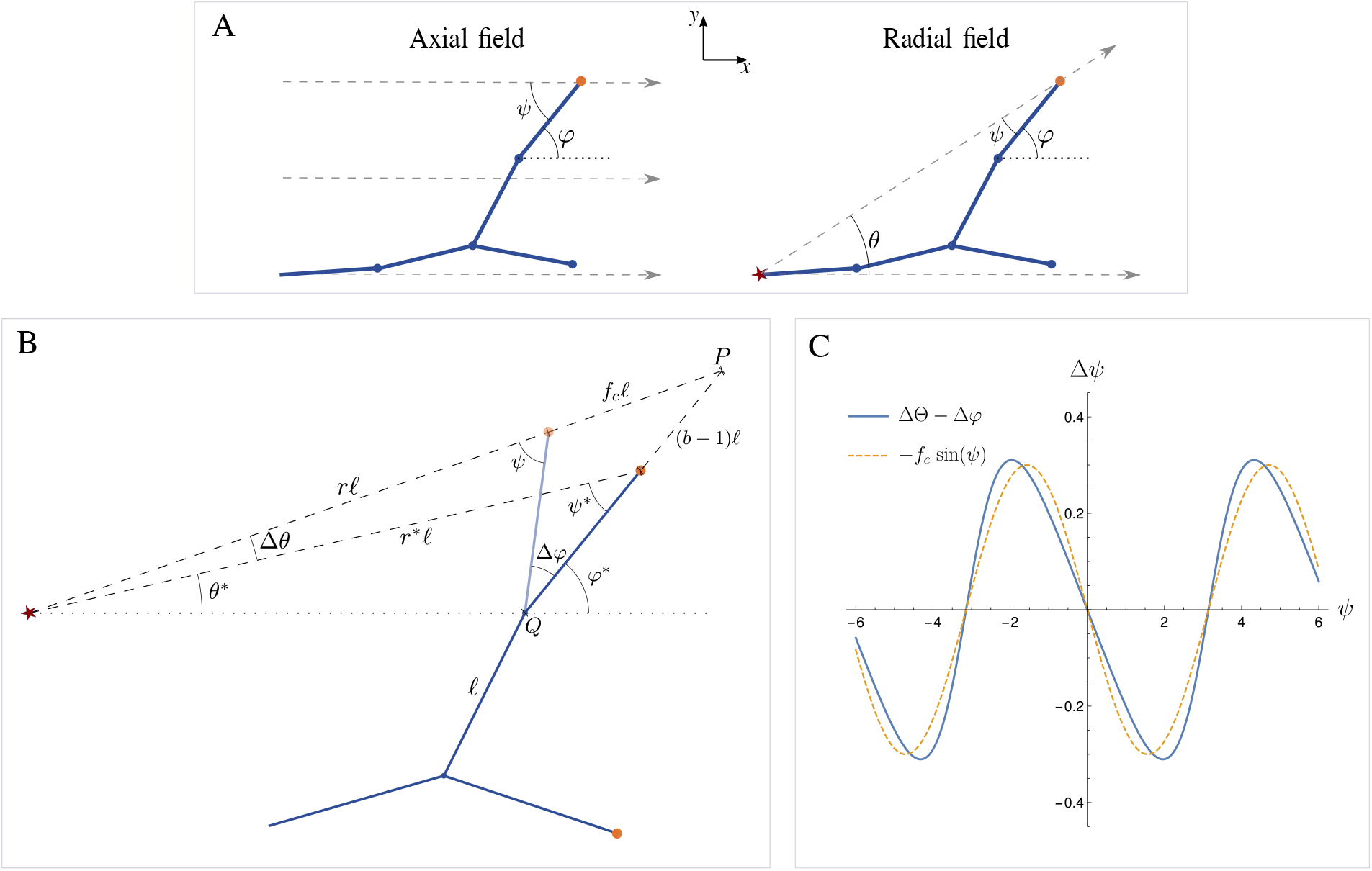
Definition and schematic of the variables used in the continuum model. (A) Schematic of branch segments in an axial (left) or radial (right) external field (dashed arrows) in two-dimensions (with *x, y* coordinates as shown) highlighting the main variables used in the analysis: In a radial field (right), for every branch segment with an active tip (orange node) one can define the *local angle φ* that it makes with the horizontal axis (dotted lines), its *angle θ to origin* (star symbol), and the *angle difference ψ* ≡ *φ* – *θ*. Accordingly, the alignment of a branch segment with the field is determined by the angle difference *ψ*, whereas for an axial field (lefty), the origin of the external field is located at *x* → −∞ (correspondsg to *θ* = 0), and thus the alignment will be determined by the local angle *φ* only (*ψ* = *φ*). Blue solid lines represent (static) branch segments of size *ℓ* generated by tip elongation (blue circles indicate inactive tips after an annihilation event due to proximity to other branches) (B) Displacement of an active tip due to a radial field. The external field acts along the radial distance *rℓ* of the active tip to the origin of the potential (labelled by star) with a strength determined by the coupling parameter *f_c_*. The branch segment is then aligned towards the extended point *P* but preserves its length *ℓ*. The angle difference after displacement *ψ*\* = *φ*\* – *θ*\* is then given by Eqs. (S2–S4), and can be approximated as a function of the field strength *f_c_* and of its value *ψ* before displacement only, see Eq. (S5). (C) Plot of the changes in the angle difference Δ*ψ* after the displacement due to the external field *f_c_*. The exact expression given by Eqs. (S2–S4) is compared with the approximation Δ*ψ* = −*f_c_* sin(*ψ*) for a value of *f_c_* = 0.3, showing good overall agreement. For smaller values of *f_c_* that are explored in the implementation of tip displacement-based simulations, the two expressions converge onto each other.

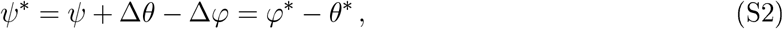

where Δ*φ* ≡ *φ* – *φ*\* and Δ*θ* ≡ *θ* – *θ*\* denote respectively the changes in the local angle and in the angle to origin values. To express the new angle *ψ*\* in tern is of *ψ* and the external field strength *f_c_*, we deduce the following trigonometric relations from simple geometric arguments, see Fig. S1(B):

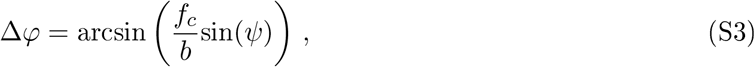

and

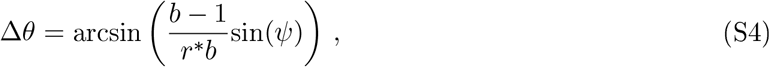

where *r*\* represents the distance of the active tip to the field origin *after* the displacement by the external field, and *b* determines the distance *bℓ* between the point *P* and the branch node *Q*, see Fig. S1(B). For a wide range of values of *f_c_*, variations in the loca 1 angle Δ*φ* strongly dominate over the term Δ*θ*, such that the latter can be neglected. Furthermore, for the weak field case that we investigate here, *i.e*., for low values of *f_c_*, the term Δ*φ* can be simply expressed by *f_c_*sin(*ψ*) because the length scale *b* takes values close to 1. The change in the angle difference after displacement can then be approximated by the following expression:

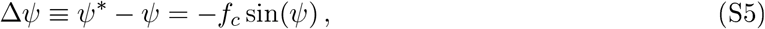

which is a function of *f_c_* and *ψ* only. Fig. S1(C) provides a comparison between this approximation and the exact relation determined by Eqs. (S2–S4). This dependency on the external field *f_c_* reflects a sinusoidal reorientation model for the angle difference *ψ* which was previously demonstrated for the gyrotaxis of spherical microswimmers [2], for cyanobacterial circadian oscillators [3] and for dipoles in a constant external field [4].

#### S1.2 Fokker-Planck equation with non-local jumps

In simulations of branching and annihilating random walks (BARWs), elongation events lead to a small rotational diffusion of the branch segments, whereas branching events lead to large, abrupt changes in the local angle values of the active tips, as illustrated in Fig. S2(A). These two different sources of noises are described by an *a priori* probability distribution λ(*φ* – *φ′*) for the difference in the local angles before *φ′* and after *φ* a jump. The external field then subsequently acts to modify these jump probabilities. This simulation setup can thus be described by separating the jumps arising from elongation and branching events from the bias arising through the external potential. Because of the large, non-local jumps in the local angle values *φ*, the standard method of reaching the continuum limit by expanding around small Δ*φ* cannot be performed. We will therefore follow a generalized method developed for Levy flights in Refs. [5, 6] to derive the Fokker-Planck equation.

We first derive the central result of Ref.[6] for generic non-local jumps in one spatial dimension (represented by the continuous variable *x*) and then apply it to the angle difference *ψ*. We start with the continuum version of the difference equation for the probability distribution *P*(*x, t* + Δ*t*) after a jump from site *x′* to *x* within the time interval Δ*t*:

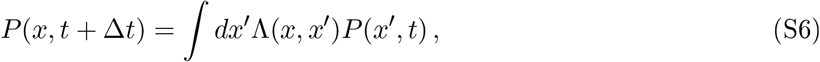

where the transfer kernel

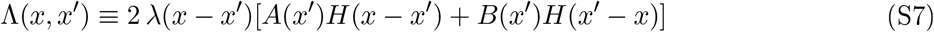

encodes both the distance between the jumps *x* – *x′* via the corresponding probability density function λ(*x* – *x′*), and the spatial dependency of the transition probabilities *A*(*x*) and *B*(*x*). *H*(*z*) denotes the Heaviside step function defined as *H*(*z*) = 1 for *z* ≥ 0 and *H*(*z*) = 0 otherwise. We furthermore assume symmetric jump distance distributions of the form λ(*x* – *x′*) = λ(*x′* – *x*), *i.e*., the probability for different jump sizes only depends on the size of the jump and not on the direction. The directional biases are governed by the transition probabilities *A*(*x*) and *B*(*x*). The normalization for the transfer kernel is defined by integrating Eq. (S7) over the jump differences *z* ≡ *x* – *x′*:

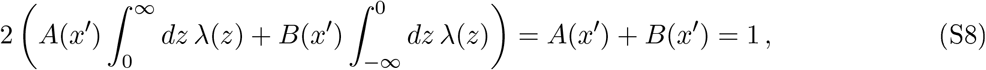

where we used the symmetry of λ(*z*) together with its normalization 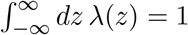. We now multiply Eq. (S6) by *e^ikx^* and integrate over *dx* to obtain

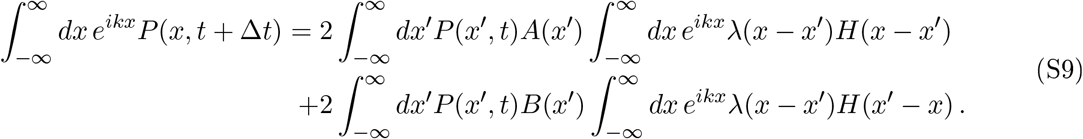

Substituting *z* = *x* – *x′* in the integrals over *x* and using the Euler identity we get:

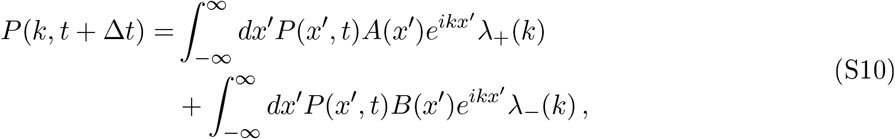

where we introduced the Fourier transform of *P*(*x, t*) and define

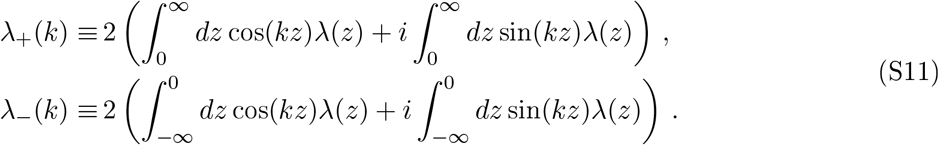

By switching the integration bounds for λ_−_(*k*) and using symmetry properties of sine and cosine functions, we can express Eq. (S10) as follows

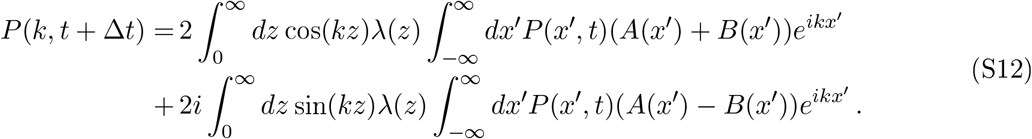

We now introduce the cosine and sine transforms

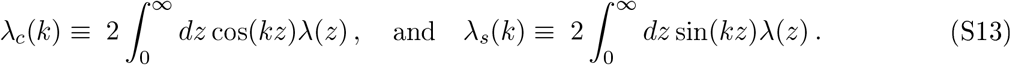

Using the convolution theorem for the second term in Eq. (S12) and recalling the relation *A*(*x*) + *B*(*x*) = 1, see Eq. (S8), we obtain the simple expression

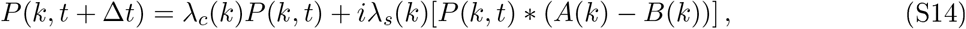

where * denotes convolution in Fourier space, as derived previously in Ref. [6].

#### S1.3 Fokker-Planck equation for the angle difference *ψ*

We now use this framework for non-local jumps to obtain an equation describing the temporal evolution of the probability distribution for the angle *ψ*. In general, changes in *ψ* after each elongation or branching event depend on the changes in the local angle *φ* – *φ′* as well as on the changes in the angle to origin *θ* – *θ′* values. However, after a small number of steps taken from the origin, the *θ*-dependent terms become negligible compared to changes in the local angle *φ*. The probability distribution for the jumps sizes of the angle difference *ψ* can therefore be approximated by that of the local angle *φ*. In fact, as we will show later, see Fig. S2(B) and (C), the mean-squared displacements for these two angles will attain the same linear form for sufficiently large times, indicating that the free diffusion of these two angular variables follow the same dynamics in this regime. As illustrated in Fig. S2(A), the jumps in the angle values are described by uniform distributions with maximal jump sizes determined by *δφ_e_* – *ψ_e_* = *π*/10 and *δφ_b_* – *ψ_b_* = *π*/2 for elongation and branching events, respectively. Denoting the branching and elongation probabilities by *p_b_* and *p_e_*, respectively, and the change in the angle difference after each jump by 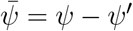, we can express the jump size distribution as

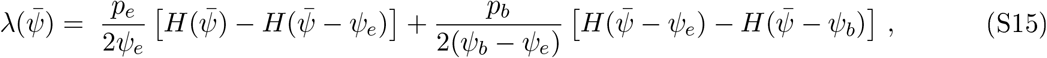

where for clarity we only express the positive part of the symmetric jump distribution, *i.e*. for 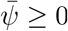. Fourier cosine and sine transforms of 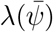 are then given by

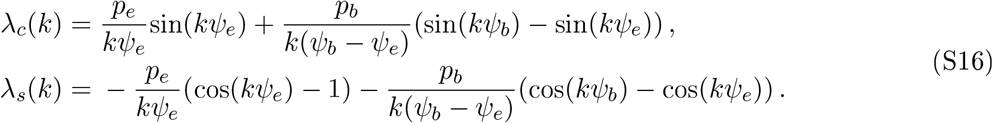

Now we use the Taylor expansions up to 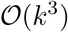 for the cosine and sine functions to obtain:

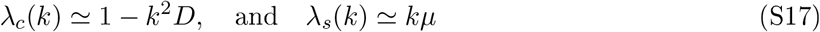

where

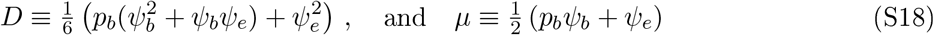

can be defined as the diffusion and mobility/advection coefficients based on microscopic variables. Inserting these expressions into Eq. (S14) and taking the inverse Fourier transform leads to

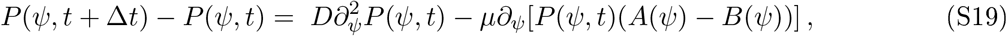

We must now define *A* and *B* to model the external field which acts to reduce the alignment angle |*ψ*| at each jump. Recalling our discussion on a radial external field, we introduce a sinusoidal force as described by Eq. (S5). Because *A*(*ψ*) + *B*(*ψ*) = 1, see Eq. (S8), and we want the drift on the particle to be determined by *A*(*ψ*) – *B*(*ψ*) = −*f_c_*sin(*ψ*), we get

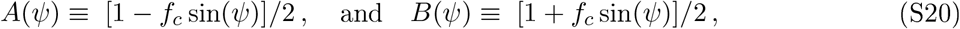

Inserting these expressions into Eq. (S19), and assuming identical mean stepping times *τ* ≡ *t/n* in the limit of large step numbers *n*, we obtain the Fokker-Planck equation

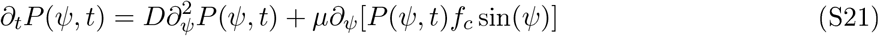

In the steady state we impose no-flux boundary conditions such that

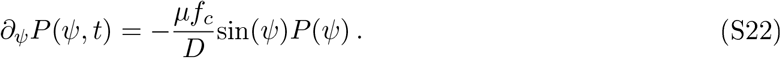

At steady state, using the normalization condition 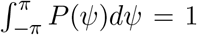 as well as the symmetry of the cosine function, this predict that *P*(*ψ, t*) should follow a von Mises distribution

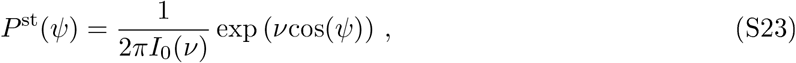

where

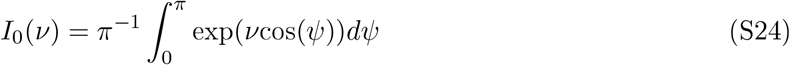

is the modified Bessel function of the first kind of order zero and

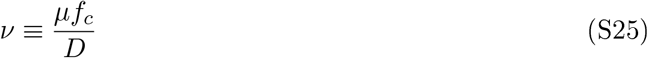

takes the role of the concentration parameter of the von Mises distribution. This is a central result of our analytical model, which we confront to experimental data in the main text. Importantly, this predicts the distribution of angles up to a single rescaled parameter *ν*, which in analogy to the Péclet number quantifies the respective contribution of advection (arising from an extrinsic guiding field) to diffusion (arising from the intrinsic stochasticity of branching/elongation events) in the Fokker-Planck equation.

Note that the variance of the angle distribution *ψ* at steady state is thus directly related to this rescaled parameter:

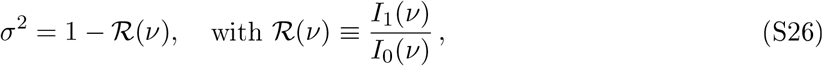

whereas the circular standard deviation takes the form [7]

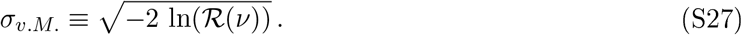

As *f_c_* → 0, this expression deviates from a power-law relationship between *f_c_* and *σ*, in contrast with the standard deviation of the equilibrium distribution for a particle in a harmonic potential, which corresponds to a normal distribution for *ψ* and depends on the external field *f_c_* as

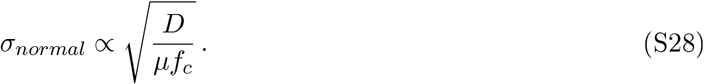

However, for sufficiently large values of *ν* that we investigate here, the von Mises SD given by Eq. (S27) exhibits a power-law behavior with a slightly larger decay coefficient, see Fig. 1 (E) for a comparison of the two scaling laws.

### S2 Simulation details

Next, we briefly summarize the details for the numerical algorithm to simulate branching and annihilating random walks (BARWs) in the absence of external field or self-avoidance.

#### S2.1 Branching and annihilating random walks (BARWs)

Similar to Ref. [8], we define active tips that will elongate and branch randomly, taking discrete steps of unit size *ℓ* = 1 per discrete time interval *τ* = 1 and leaving behing inactive branch segments that remain immobile at coordinate **r** ≡ (*x, y*) on a two-dimensional plane. When an active tip comes in close proximity of an inactive branch segment, *i.e*., when the inactive branch is within an annihilation radius *R_a_* of the active tip, the latter will become inactive and immobile. We choose a rather small annihilation radius of *R_a_* = 1.5*ℓ* to mimic contact-dependent membrane recognition. Note that the specific choice of the annihilation radius in two dimensions does not influence the overall topology (up to global rescaling) of the networks because intersection of two branches occurs with probability 1 for all sufficiently small radii. In the case of neurons, as discussed in the main text, this radius could be effectively higher to take into account the cycles of tip growth and retraction when encountering neighboring branches. Coarsening this in the model by running simulations for larger *R_a_* = 8*ℓ* (see next section) did not qualitatively change the results for small self-repulsion, but for large *f_s_*, the angle distributions became markedly sharper with faster decaying tails than the von Mises scaling-a case we did not observe in experimental data.

Simulations start with a single active tip at an initial position **r**_0_ ≡ **r**(*τ* = 0) = (0, *y*_0_) with a pre-defined polarity **p**_0_ ≡ (1, 0), *i.e*., which is directed horizontally towards the right (higher *x* values), and proceed until a certain maximal time *τ*_max_ = 200*τ* is reached. Because the active tips take a single step at each time interval *τ*, the maximal time *τ*_max_ of the simulation also determines the maximal distance *R*_max_ from the origin **r**_0_ that the last surviving active tips can attain. For the experimental data, this parameter is therefore strongly constrained by the geometrical properties (overall size and shape) of the fin, which we determine by counting the *maximal* number of steps in the coarse-grained reconstruction of the filaments, see section S3.5.2. The initial polarity of the starting active tip allows us to define an initial *local angle φ*_0_ = 0 with respect to the *x*-axis associated with the tip. The key parameter in this simulation setup is the branching probability *p_b_*, which will determine the frequency of branching events and thus also the final density of the network.

The (i) elongation and (ii) branching events for the active tips are implemented in the simulation as follows: At each time point *t*, we draw *n_a_* (pseudo-)random numbers *r_j_* ∈ (0,1) with *j* = 1,…, *n_a_*, where *n_a_* is the number of active tips at *t*. For each active tip, the elongation or branching occurs for *r_j_* > *p_b_* or *r_j_* ≤ *p_b_*, respectively. (i) When an active tip at coordinate **r** with local angle *φ* undergoes an elongation event, it takes one step “forward” such that its new local angle *φ′* will take a random value uniformly distributed between *φ* ± *δφ_e_* with *δφ_e_* = *π*/10. The latter rule leads to a small rotational diffusion of the tip during elongation events. The new position of the active tip will then be given by **r′** = **r** + (*ℓ*cos(*φ′*), *ℓ*sin(*φ′*)). Note that for *δφ_e_* = 0 the rotational diffusion vanishes and the random walk becomes infinitely persistent, (ii) When an active tip with local angle *φ* undergoes a branching event, it will produce two new active tips at positions 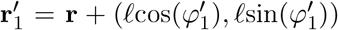 and 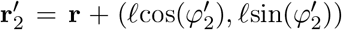, respectively, and will become inactive itself. The local angles 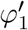 and 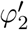 of the two new active tips respectively take values uniformly distributed in [*φ* + *δφ_e, φ_* + *δφ_b_*] and [*φ* – *δφ_e, φ_* – *δφ_b_*] with *δφ_b_* = *π*/2, *i.e*., the two new tips will be located on the two different sides of the line determined by the polarity vector **p** = (*ℓ*cos(*φ*), *ℓ*sin(*φ*)) of their parent while preserving a minimal angle of 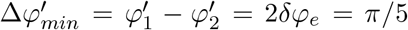 between each other. The latter rule enforces a minimal distance between the two new tips to reduce the frequency of immediate annihilation of the two new tips. Fig. S2(A) illustrates the elementary steps implemented in the simulation for the branching, elongation and annihilation of the active tips.

To test our analytical predictions for the “free” diffusion of branch segments, we first ran simulations for single branches that perform elongation and branching jumps without generating additional new active tips. We found that the mean-squared displacements of the final local angle *φ* of single branches at time *t*_max_ closely follow the relation 〈Δ*φ*(*t*_max_)^2^〉 = 2*Dt*_max_ with the diffusion constant *D* given by the “microscopic” expression Eq. (S18), see Fig. S2(B). We also obtained the same relation for the mean-squared displacements of the *angle differences, i.e*., 〈Δ*ψ*(*t*_max_)^2^〉 = 2*Dt*_max_ for large values of *t*_max_, see Fig. S2(C). The latter results justified our hypothesis that for sufficiently large times, the instantaneous jumps in the angle difference values *ψ* are dominated by the changes in the local angle *φ*, and thus do not depend on the angle *θ* to origin.

For the full BARW simulations, we could then set a branching probability *p_b_*, the initial position **r**_0_, and the initial polarity vector **p**_0_, and produced branching networks that radially grow in all directions over time with active tips forming a front, leaving inactive branches of constant density in the “inner” regions. When the network growth is constrained within a spatial region delineated by fixed boundaries, we observe an apparent directionality as predicted previously [8], see Fig. S3(A). However, tissue growth in the absence of spatial boundaries always remains isotropic, see Fig. S3(B), which also remains valid for branching networks with strong self-avoidance. We concluded that one needs to define an external field to guide the tips for a directed (anisotropic) spatial invasion such as seen in the experiments.

#### S2.2 BARWs with external guidance and/or self-avoidance

In this subsection, we describe different possible microscopic ways to implement external guidance in the simulations of branching and annihilating random walks, and show importantly that the two give rise to the same qualitative predictions in the continuum model in terms of overall branch alignment.

**Figure S2:**
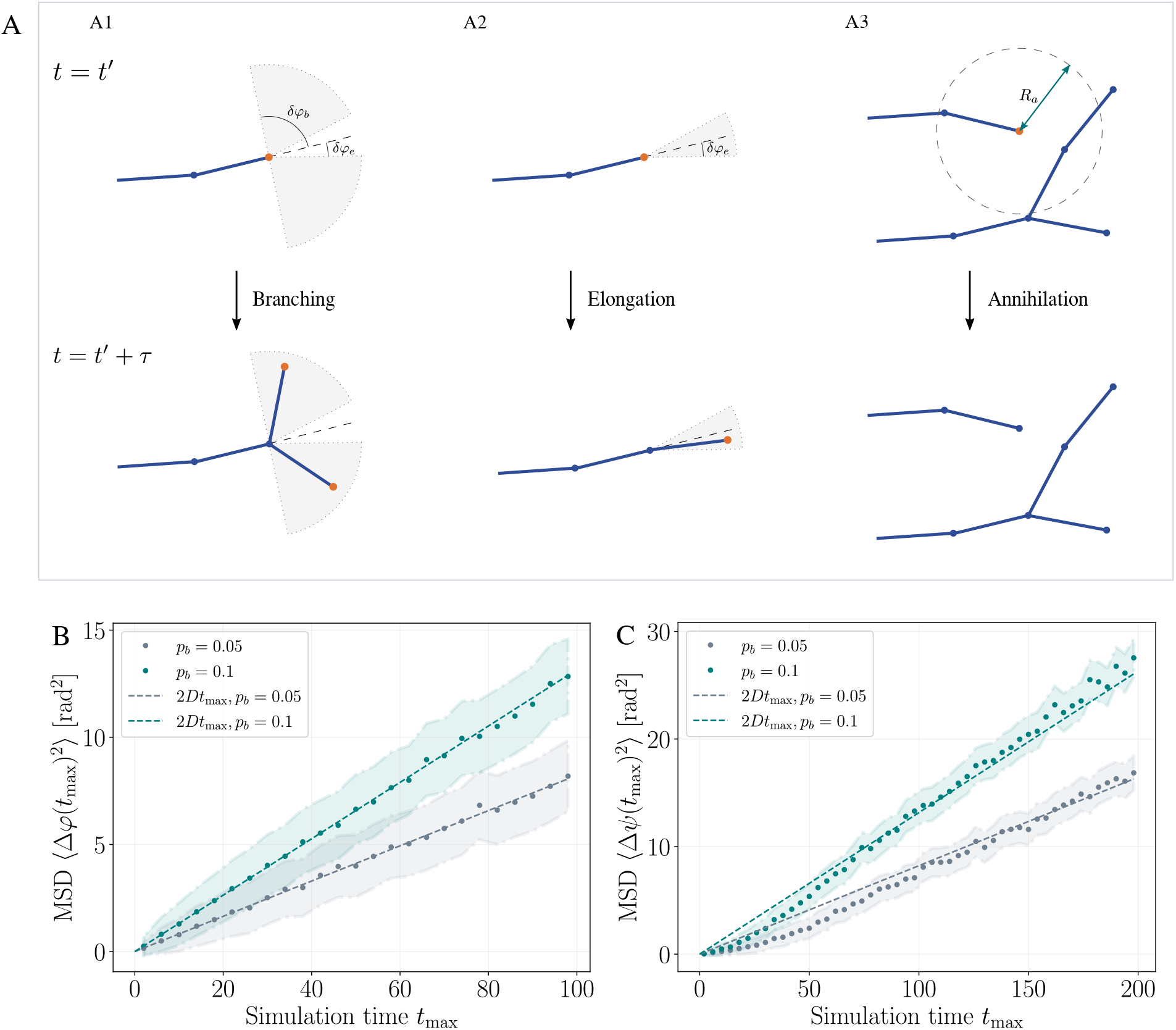
(A) Elementary steps in the implementation of the BARW simulations: (A1) With a given *a priori* branching probability *p_b_*, an active tip (orange node) of a branch segment at time *t′* generates two additional active tips at the next time step *t′* + *τ*, where *τ* is a discrete time interval (set to *τ* = 1 in the simulations). The angles between the new branch segments with the previous segment take values uniformly distributed in ±[*δφ_e_, δφ_b_*] (gray shaded areas), where we use *δφ_e_* = *π*/10 and *δφ_b_* = *π*/2. (A2) Elongation of an existing branch segment occurs with the probability *p_e_* = 1 – *p_b_* and leads to a small rotational diffusion of the branch confined within a cone determined by the angle 2*δφ_e_* (gray shaded area). (A3) If an active tip comes in close proximity of existing inactive branch segment (blue nodes) it will annihilate, *i.e*., become inactive. The local neighborhood (dashed circle) for sensing inactive branches is determined by an annihilation radius *R_a_* (green arrow). (B) Simulated trajectories of a single branch segment freely diffusing without external potential and annihilation displays a mean-squared displacement (MSD) for the local angle *φ* that obeys the relation 〈Δ*φ*(*t*_max_)^2^〉 = 2*Dt*_max_ with the diffusion constant *D* predicted by the microscopic theory, see Eq. (S18). (C) MSD for the angle difference *ψ* of a single branch segment approaches the analytical prediction for large times (*t*_max_ ≥ 100). Plot markers represent the mean values and the shaded regions denote circular SDs around the mean values (*n* = 1000 and *n* = 2000 runs for (B) and (C), respectively).

##### S2.2.1 Microscopic mechanisms for implementing external guidance

Observations of the movement of single neuronal tips in the presence of chemical cues (such as Netrin) indicate that the neuronal tips can reorient themselves even without neighboring tips or other branches in close proximity [9]. This change in the directionality can be effectively modelled as a displacement of the tip towards the external field, as we discussed in section S1.1.1 and represent in Fig. S1. Note that, however, such a displacement would lead to an instantaneous shift (jump) in the angle difference value *ψ* of the active tip due to the external field in contrast with the the continuum model developed in section S1.3.1, according to which the external field influences the *jump probabilities* for the angle difference *ψ via* the forward and backward bias terms defined in Eq. (S20). We will therefore discuss these two different ways of implementing the external field, and show that they generically result in the same behavior and the angular alignments follow the same statistics up to a constant prefactor for the field strength.

**Figure S3:**
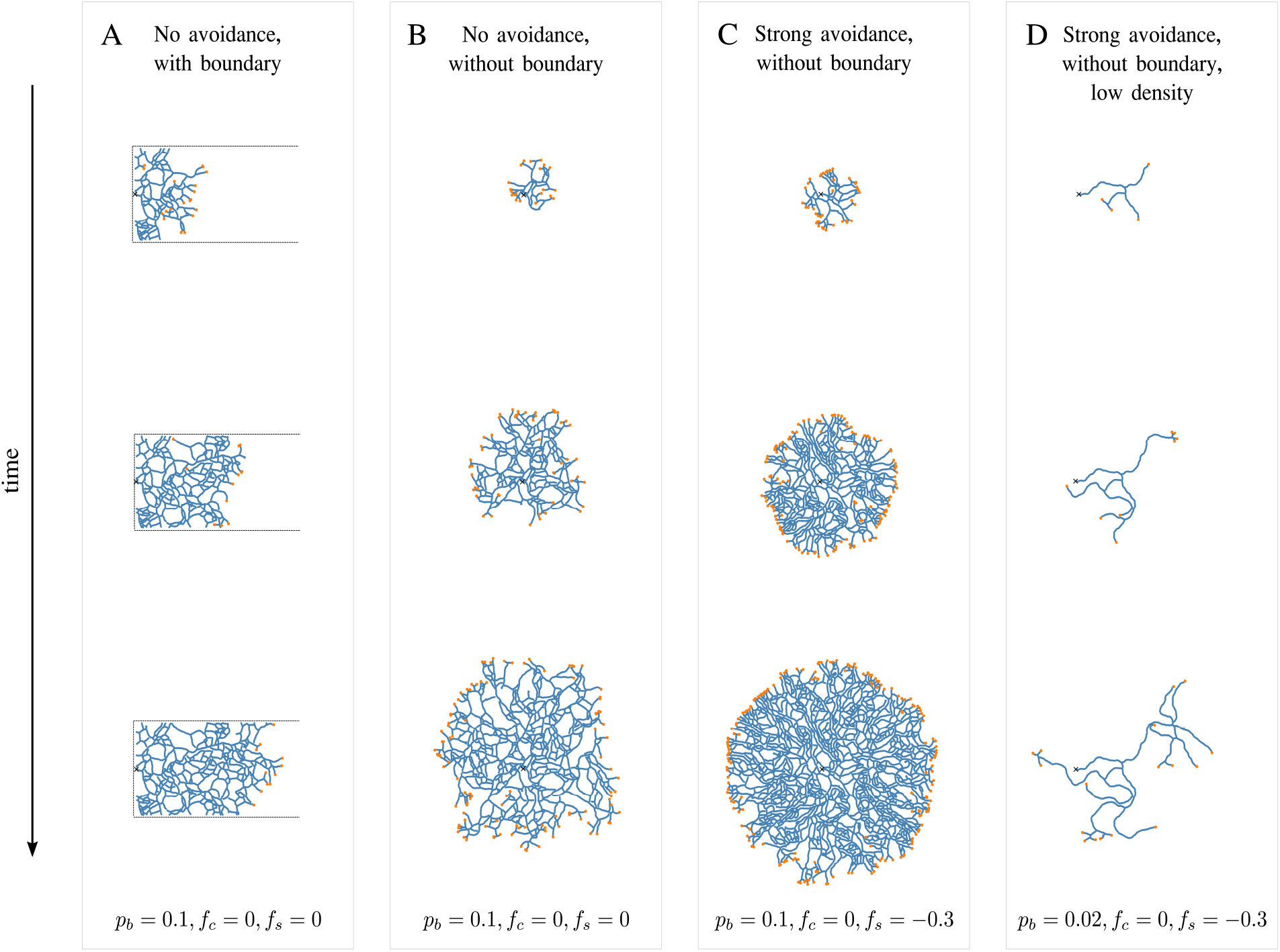
Time evolution of BARWs in the absence of an external field (*f_c_* = 0). All networks start growing from the initial point denoted by the cross symbol (black). (A) Simulation of BARWs confined in a bounded region (represented by the dashed rectangular half-open box) exhibits directed growth of the branched network effectively towards the opening on the right. (B) When the boundary is removed, the branching network grows isotropically in all directions, where active tips leave passive branch segments in the inner regions of the network. (C) BARWs with a rather strong self-avoidance (*f_s_* = −0.3) generate a dense network with efficient space-filling (as described in Fig. 5 of the main text), where active tips seem to define a sharp front that propagates isotropically. (D) Reducing the branching probability results in a network with a small number of active tips and a well-defined front does not emerge. Despite the strong self-avoidance, an anisotropic growth cannot be observed in this purely local, self-organized model in the absence of external field.

###### External field *via* biased probabilities

To closely follow the continuum model, we first implemented biased jump probabilities in the simulation as follows: For each elongation event, we weighted the elongation probability *p_e_* = 1 – *p_b_* by the bias terms such that a “forward” and “backward” step that respectively increases or decreases the angle difference value *ψ* occurs with the probability *p_e_ A*(*ψ*) and *p_e_ B*(*ψ*) with *A*(*ψ*) = [1 – *f_c_*sin(*ψ*)]/2 and *B*(*ψ*) = [1 + *f_c_*sin(*ψ*)]/2 as previously defined in Eq. (S20). Here, *f_c_* denotes the strength of the external field. Because *A*(*ψ*) + *B*(*ψ*) = 1, the joint probability of a forward and backward elongation event is equal to *p_e_* which is the *a priori* value fixed by setting the branching probability *p_b_*. For instance, for an active tip with a positive angle difference value 0 < *ψ* < *π*, this implementation of the biased jumps implies that it is more likely for the tip to reduce its local angle than to increase it upon elongation. For branching events we do not implement the effect of the external field *f_c_* on the angle values of the tip because the bifurcation event in the simulation corresponds to two opposite jumps in the local angle values (the two tips branch into different sides of the polarity vector). In contrast, the jumps corresponding both to elongation and branching events in the continuum model are influenced by the external field *via* the bias terms *A*(*ψ*) and *B*(*ψ*). Even though we omit the bias for the branching events, we obtained a good agreement with the analytical predictions for the angle distributions obtained using this simulation setup, as shown in the main text.

###### External field *via* tip displacement

We also wanted to explore the modelling of the external field in a different way, to show the generality of our approach and the insensitivity to the details of the microscopic tip behavior, and thus implemented guidance *via* tip displacements. In this simulation setup, the active tips would now be displaced at each time step by a factor *f_c_* **p**_*c*_, where **p**_*c*_ ≡ (*ℓ*cos(*θ*), *ℓ*sin(*θ*)) is the unit vector pointing along the fie Id lines, and *θ* denotes the angle to origin of the tip before displacement. In this second case, the field thus directly “corrected” the directionality of the tip migration, as opposed to the “effective correction” which arose from the biased transition probabilities of the previous approach. Even though these two ways of implementing the external field are quite different, we found that they generated both qualitatively and quantitatively very similar network structures, see section S2.4 for a comparison of the results. Crucially, we observed that one only needs to tune the external field strength *f_c_* by a constant prefactor *α* to obtain similar results using the two different algorithms.

##### S2.2.2 Implementing self-avoidance

An additional feature observed in some branching tissues is the presence of self-avoidance of growing tips, *e.g*., as clearly evidenced in morphogenesis of starburst amacrine cells [10], where active tips sense, and are repulsed by, existing branch segments in close proximity. For neuronal branching it was shown that this self-avoidance of the growing dendrites is based on isoform recognition of the membrane proteins [11]. We modelled this effect by considering that active tips sense an average density vector **p**_*s*_ [8] depending on the number of branch segments within a certain radius *R_s_* of “self-recognition”:

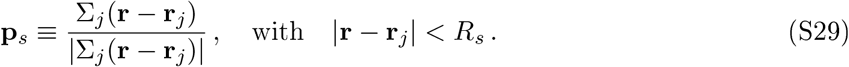

Here, for notational simplicity, we denote the position of the active tip by **r** and that of the remaining particles within *R_s_* by **r**_*j*_. The density vector **p**_*s*_ will thus define a normalized vector pointing *away* from nearby particles. We can now define a self-recognition force by – *f_s_***p**_*s*_, where *f_s_* determines the strength of interaction with *f_s_* > 0 and *f_s_* < 0 corresponding to attraction and repulsion, respectively. Note that this self-repulsion force can effectively correspond to sequential recognition vs. retraction events such as seen in neuronal dendrites and as modelled *e.g*. in Ref. [12]. In contrast with the latter study, here we do not focus on the statistics of the retraction events and could therefore describe the self-recognition *via* a more simplified effective modelling. The position of the active tip after displacement by this self-recognition force will then simply change to **r**\* = **r** – *f_s_***p**_*s*_. However, in order to preserve the step length *ℓ*, we will also correct the final position such that the distance between the final position **r**\* after displacement and the position *at the previous time step* is equal to *ℓ*, *i.e*., |**r**\*(*t*) – **r**(*t* – *τ*)| = *ℓ*.

**Figure S4:**
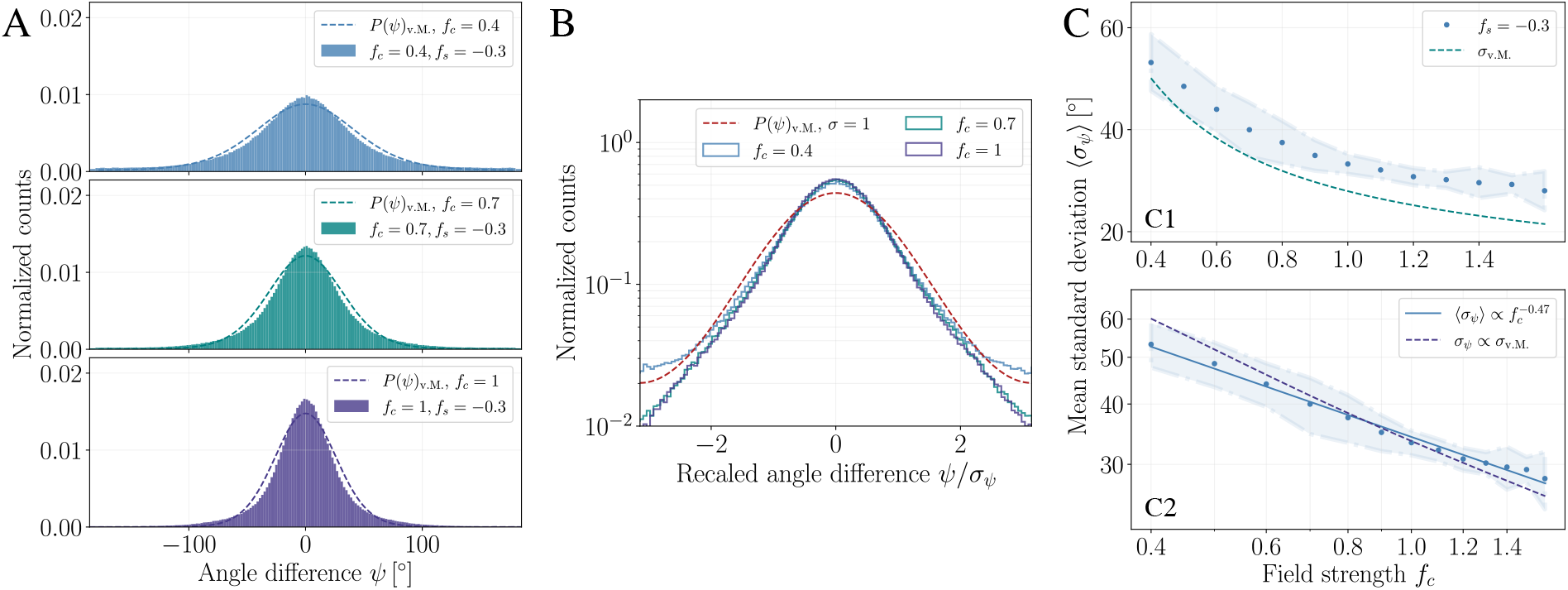
Effects of strong self-avoidance (*f_s_* = −0.3) on the angular alignment. (A) Normalized histograms for the angle difference *ψ* for different values of the external field strength *f_c_*. Dashed lines represent the theoretical predictions given by the von Mises distribution with the concentration parameter 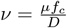. Presence of strong self-avoidance leads to narrower distributions for *ψ* and to a strong deviation from the analytical prediction for large *f_c_*, in contrast with the case without self-avoidance, see Fig. 3B in the main text. (B) Normalized histograms for *ψ* rescued by their corresponding mean standard deviations (SDs) *σ_ψ_* (solid lines) compared to the von Mises distribution with unit SD (dascd line) exhibit a sharper peak and pronounced deviations at the tails for large *f_c_*. (C) Mean SDs of the normalized histograms for *ψ* decay monotonically as a function of *f_c_* and attain larger values than the SDs corresponding to the von Mises distribution (dashed line, C1). (C2) Scaling of the SDs obeys a power-law 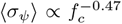 close to the scaling precited by the von Mises distribution (dashed line).

#### S2.3 Simulation results for BARWs with self-avoidance

##### S2.3.1 Strong self-avoidance without external field

To clarify whether directed growth can arise in a self-organized way in the absence of an external field just by the self-avoidance mechanism, we performed simulations with a strong self-avoidance of *f_s_* = −0.3 but setting the external field strength to *f_c_* = 0. In the density regime set by a branching probability of *p_b_* = 0.1, we again observed isotropic growth of the networks, see Fig. S3(C). However, due to the the rather strong self-avoidance potential, the final network exhibited a qualitatively much “ordered” structure, in agreement with the predictions in [8]. Interestingly, the active tips formed a sharper propagating front in contrast to the case without self-avoidance. We finally analyzed the low-density case by reducing the branching probability to *p_b_* = 0.02 to see if the network could preserve its pre-defined directionality uninterrupted by the “outwards push” due to branching events. However, such a self-organized anisotropic growth also could not be obtained for the low-density case, see Fig. S3(D), which indicated that an external field is required to break the isotropy in the tissue growth.

##### S2.3.2 Strong self-avoidance with external field

Here we provide further results for simulations with a high self-avoidance potential and in the presence of external guidance. For increasing values of *f_s_*, we observed that the density and the space-filling properties of the network markedly increased, as shown in Fig. 5 in the main text. We also qualitatively observed that the networks with high self-avoidance exhibit better alignment with the external field. This increased alignment is indeed reflected in the angle distributions, where the distributions for the angle difference become narrower compared with those from simulations without self-avoidance, see Fig. S4. The mechanism can presumably be linked to the high density of the network and its space-filling properties, where self-avoidance competes against annihilation events, thus the network effectively generates more segments in dense regions that are forced to be aligned with the external field. Supporting this hypothesis further, we see faster decaying tails for large values of *f_c_* and for strong-repulsion, as exhibited by the rescaled angle difference histograms in Fig. S4(B).

**Figure S5:**
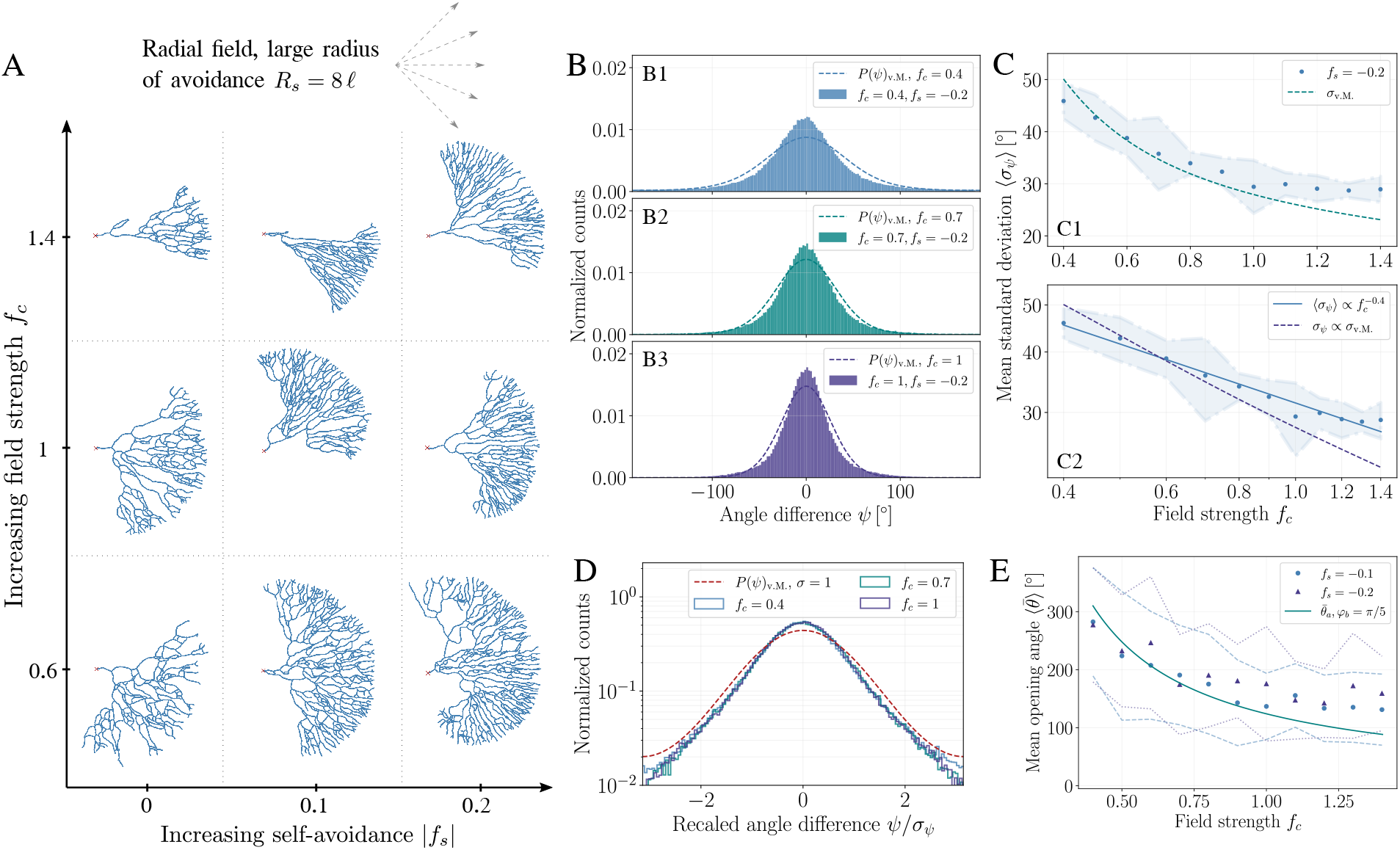
Effects of a large radius of self-repulsion (*R_s_* = 8*ℓ*) on the alignment and morphology of branching networks. (A) Morphology diagram of branching networks in a radial field. For strong self-repulsion, individual branches exhibit pronounced alignment with characteristic gaps larger than those seen for smaller radius of avoidance as used in the main text (see Fig. 2A in the main text with *R_s_* = 3*ℓ*). (B) Normalized histograms for the angle difference *ψ* for *f_s_* = −0.2 display sharper peaks than those of the von Mises distributions predicted by theory (dashed lines). (C1) Mean standard deviations (SDs) for *ψ* are close to theoretically predicted values (dashed line) but their scaling with increasing field strength *f_c_* deviates strongly from the scaling relation for the SD *σ*_v.M._ of the von Mises distribution (C2). (D) Histograms for *ψ* rescaled by their SDs (solid lines) illustrate sharper peaks and faster decaying tails compared to the von Mises distribution with unit SD (dashed line). (E) Mean opening angles 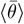 for branching networks with a large radius of repulsion for a repulsion strength of *f_s_* = −0.1 (circular markers) and *f_s_* = −0.2 (triangular markers) as a function of the field strength *f_c_*. For large *f_c_*, the opening angles saturate and deviate from the predicted decay 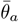 (solid line), in contrast with the case for a small radius of avoidance (compare Fig. 4 in the main text).

##### S2.3.3 Large radius of self-avoidance

To explore the alternative of self-interactions that are controlled at larger distances (for instance e.g. longer retraction events after contact-mediated self-recognition) we set a large radius of self-recognition *R_s_* = 8*ℓ* as compared with the radius *R_s_* = 3*ℓ* we used otherwise. Interestingly, even on a visual level, we could observe strongly aligned branches in the morphology diagram, see Fig. S5(A), which became quite pronounced for a large value of self-repulsion (*f_s_* = −0.2, rightmost column). Analysis of the angle distributions revealed that this case indeed leads to large deviations from the analytical predictions, and the standard deviations decay also rather slowly with increasing *f_c_*, see Fig. S5(B) and (C). For large *f_c_*, these effects of self-repulsion become rather dominant and lead to a saturation of the mean opening angles 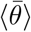 of the networks as shown in Fig. S5(E). Together, these results indicate that such long-range self-interaction effects can indeed influence the angular alignment and morphology of the final networks, especially in the presence of strong external potentials (although our data indicates that this it not the experimentally relevant limit, see Fig. 5 of the main text, this would be an interesting metric to test in other branched organs).

**Figure S6:**
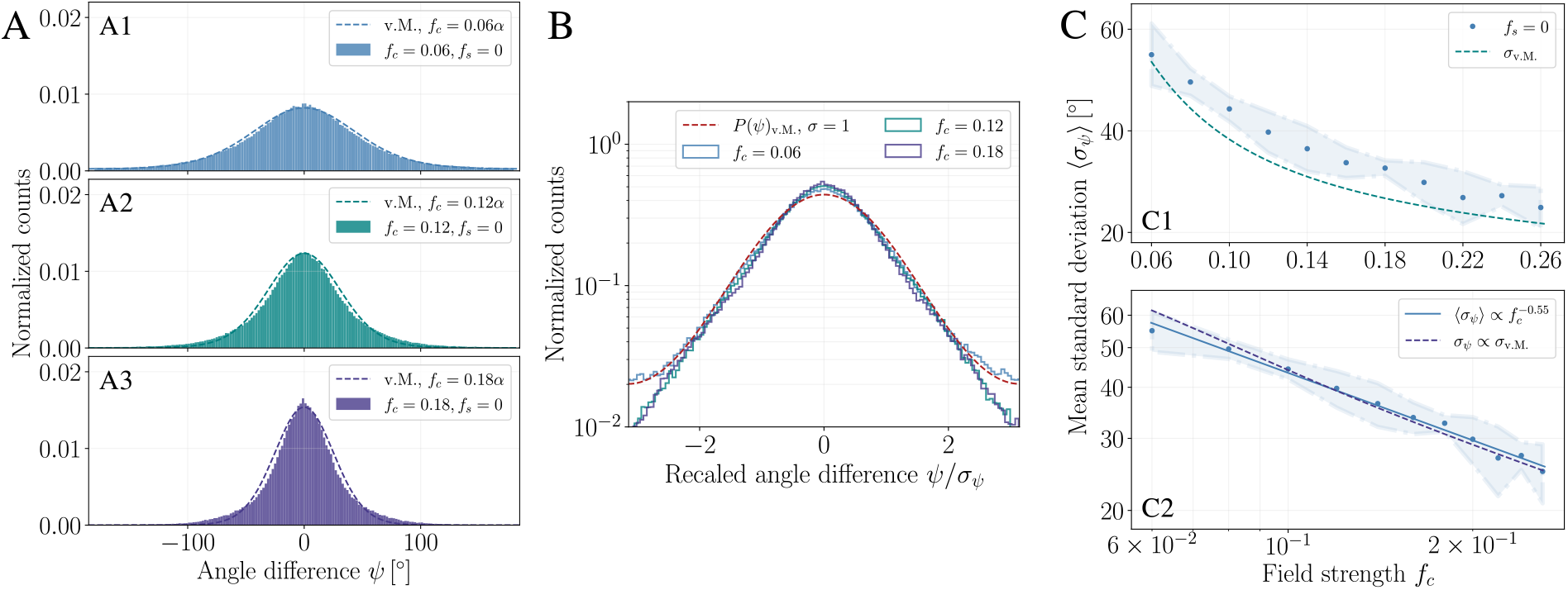
Statistics of BARWs with external guidance *via*, deterministic displacement of active tips. (A) Normalized histograms for the angle difference *ψ* for different values of the external field strength *f_c_* are well-approximated by von Mises distributions (dashed lines) with a concentration parameter 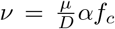 with an additional prefactor *α* = 6 used as a fit parameter. (B) Histograms for *ψ* rescaled by their corresponding standard deviations (SDs) are well-described by the von Mises distribution with unit SD (dashed line). For large *f_c_* the tails of rescaled histograms (solid lines) decay faster than that of the von Mises distribution. (C) Mean SDs 〈*σ_ψ_*〉 of the angle difference distributions decay monotonically with *f_c_* (C1) with values close to the analytical prediction (dashed line), and exhibit a power-law relation 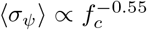 (C2) well-approximated by the analytical relation *σ*_v.M._ corresponding to the von Mises distribution (dashed line).

#### S2.4 Simulation of BARWs with external guidance *via* tip displacement

Next, we performed simulations with the alternative implementation of the external guidance *via* direct tip displacements, as we briefly described in section S2.2. First, we qualitatively observed a high similarity between the final network topologies generated using the two methods for different values of the external field strength. We then quantified this similarity by analyzing the angle distributions and the scaling of the alignment fluctuations, see Fig. S6. We found that the two methods generically provide the same statistics after tuning the field strength *f_c_* by a constant prefactor *α*. Note that for different choices of the branching probability, this pre-factor needed to be changed slightly but recovered the statistics for different *f_c_* values. To probe the relevance of direct tip displacement-based simulations, live-imaging dataset would be needed to investigate more systematically the microscopic mechanisms of tip elongation, repulsion and guidance.

#### S2.5 Estimating the average opening angle

As briefly discussed in the main text, the area of invasion of a branching network in a radial external field will be proportional to the average opening angle 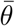 between the outermost branch segments of the network. To quantify this, we conjectured that the overall opening angle is very likely to be dependent on the local dynamics *at the growing boundary* of a branching network. Indeed, using again simple geometric arguments for the displacement of active tips by an external field, similar to our analysis in section S1.1.1, we infer

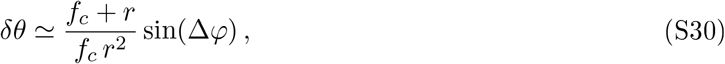

where *δθ* ≡ *θ*(*t*) – *θ*(*t* – *τ*) denotes the changes in the angle to origin values of the active tip between two consecutive time intervals, Δ*φ* ≡ *φ* – *φ*\* is the change in the local angle after the displacement by the external field *f_c_* at time *t*, and *r*(*t*) ≃ *r*(*t* – *τ*) ≡ *r* is the radial distance of the active tip from the field origin. For the dynamics “orthogonal” to the growth direction that we are interested here, the changes in the radial distance between consecutive time steps can be neglected and thus described effectively by a single parameter *r* replacing the temporal parameter *t*. After these simplifying assumptions (and assuming *δθ/dr* is a well-defined infinitesimal), we can integrate Eq. (S30) to obtain

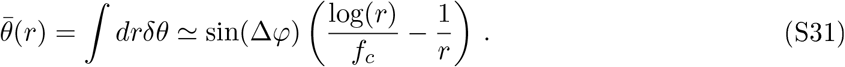

Arguing that Δ*φ* is likely to be proportional to the branching probability *p_b_* (because at large time scales the branching events simply will act to gradually increase the local angle values *φ*), we can approximate the prefactor sin(Δ*φ*) ≃ *p_b_φ_b_* with a free parameter *φ_b_*. For large times, the second term in Eq. (S31) inversely proportional to *r* becomes smaller and thus we arrive at Eq. (3) of the main text.

### S3 Experimental model system and methods

#### S3.1 Zebrafish transgenic lines and husbandry

Zebrafish were raised and housed in the Karolinska Institutet core facility following established and approved procedures. The study was performed in accordance with local guidelines and regulations and approved by Stockholms djurförsöksctiska nämiid. The new transgenic zebrafish strain was generated by injecting UAS:mCherry-caax to Tg(HuC:Gal4; UAS:synaptophysin-GFP) as described below. The resulting FO transgenic fish express red fluorescent reporter mCherry in cell membranes of a sparse number of neurons, allowing visualization and analysis of neuronal arborization *in vivo*.

#### S3.2 Cloning

The expression construct of UAS:mCherry-caax was generated with tol2 kit [13] by recombining p5E-UAS (tol2 kit // 327), pME-mCherryCAAX (tol2 kit // 550), p3E-polyA (tol2 kit // 302), and pDest-Tol2pA2 (tol2 kit // 394). The niRNA of alpha-bungarotoxin was prepared using Addgene plasmid, // 69542 as a template and niRNA of pCS2FA-transposase using tol2 kit // 396 as a template [14]; *in vitro* transcription was performed with mMcssagc niMachine SP6 kit (Thermo Fisher Scientific) and RNA was purified with RNcasy Mini Kit (Qiagen). Zebrafish embryos of Tg(Huc:gal4VP16;UAS:synaptophysin-GFP) were injected with 90 pg of alpha-bungarotoxin niRNA with 10% phenol red and 0.13 M KC1 into yolk at one cell stage. Then 20 pg of UAS:mCherry-caax and 20 pg of transposase niRNA were injected with 10% phenol red and 0.13 M KC1 into one of the cells at 4-8 cell stage.

#### S3.3 Immunostaining

For the whole-mount imaging, we anesthetized fish at the 24 hpf, 48 hpf and 5 dpf stages with Tricaine in the same manner as described above, followed by fixation with i% PFA for 4h at room temperature. Subsequently, the specimens were pcrmcabilizcd with three 30 min washes in 100% methanol, washed with PBS supplemented with 0.1%, Tween 20 (PBST) five times for 15 min, stained with the primary antibodies in blocking solution (5% normal donkey serum, 10%, DMSO, 0.1%, Tween-20, in PBS) for 48h, washed 5 times in PBST for 30 min, stained with secondary antibodies for 24h, washed in PBST as described above, and finally dehydrated in 100% methanol with two 30 min washes and rendered transparent with clearing solution consisting of one part benzyl alcohol and two parts of benzyl benzoate (BABB). The primary antibodies utilized were anti-acetylated tubulin (Neuronal marker, Gene Tex), anti-HuC/HuD neuronal protein and (Abcam), all diluted 1/800 in blocking solution. Alexa fluor 555 donkey anti-rabbit and Alexa fluor 647 donkey anti-mouse (all from Invitrogen) were used as secondary antibodies at a dilution of 1/1000 in blocking solution. Note that tubulin staining was punctiform at high magnification therefore only fish of the mCherry line were used for 3D reconstruction.

**Figure S7:**
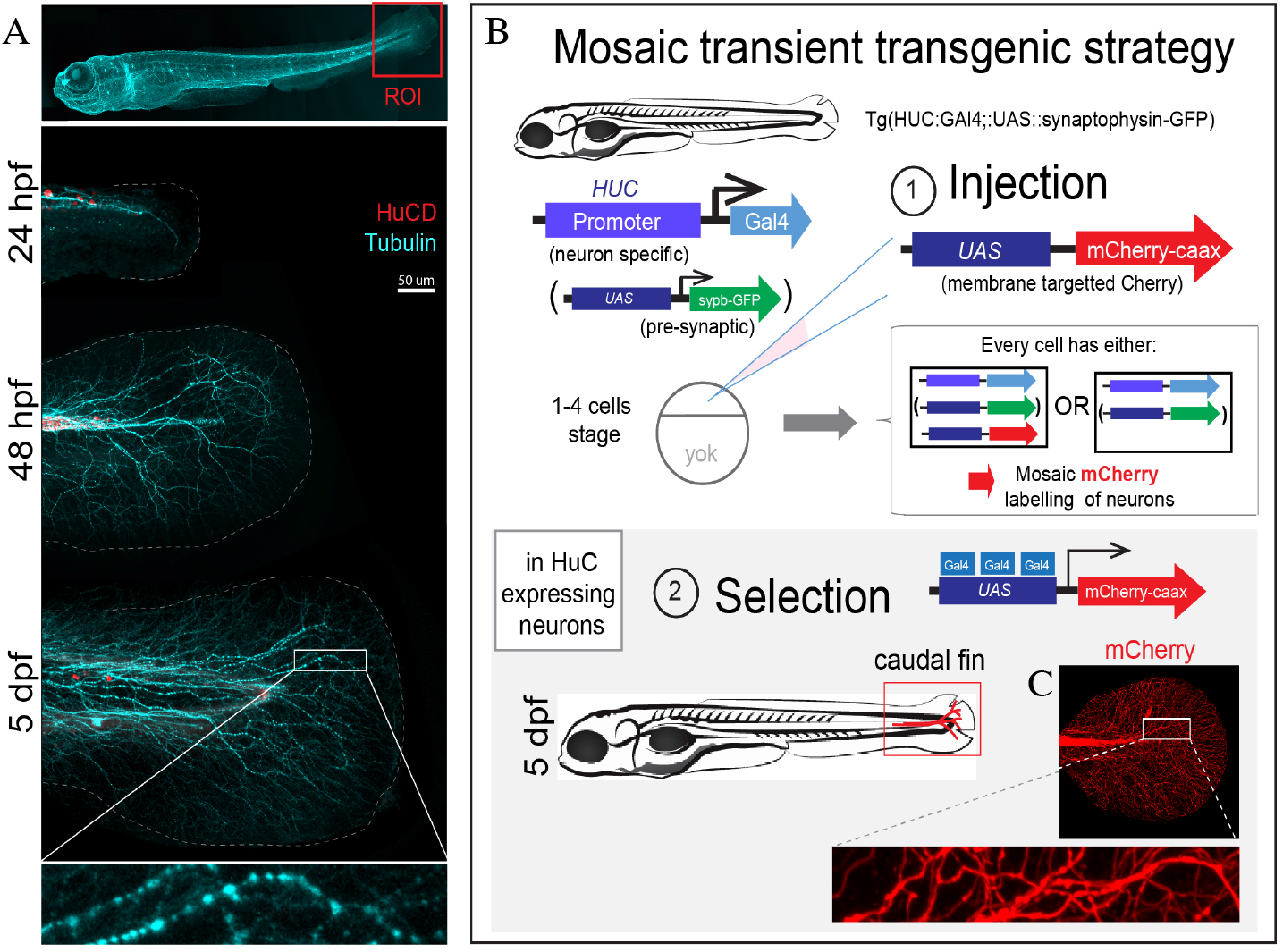
Developmental stages of caudal fin innervation, schematic of the mCherry-based experimental strategy and *in vivo* caudal fin imaging. (A) Developmental stages of zebrafish caudal fin innervation. Top panel - 5 dpf zebrafish wholemount with red box indicating the Region of Interest (ROI): caudal fin. Several other features such as neuromasts and lateral line are visible. All tubulin positive fibers are stained with anti-acetylated tubulin antibody and presented in cyan and HuC positive cell bodies are presented in red. Different developmental stages are presented −24 hpf, 48 hpf and 5 dpf. First sensory innervation starts to appear in the caudal fin area at 24 hpf, developing into a dense network of innervation by 48 hpf (egg hatching) and reaches maximum density at 5 dpf (swimming behavior). HuC/D positive cell bodies and Tubulin positive neuronal projections are visible on all stages. Caudal fin outline is marked with the dashed lines. Magnified inset on 5 dpf stage reveals the puncta-like tubulin immunostaining which compromised potential 3D reconstruction of single neuron peripheral arborization. Indeed, a continuous signal would be preferable for reliable reconstructions, as well as sparse labelling of neurons, justifying our mCherry-based strategy. (B) Experimental scheme of our Mosaic transient transgenic strategy that lead to mCherry mosaic labelling of neurons in 5 dpf zebrafish. (1) Using GAL4-Upstream Activating Sequence (GAL4-UAS) methodology, we generated transient transgenic fish in which a sparse number of neurons will express mCherry. Injection of genetic vector carrying UAS-mCherry-caax construction to 1-4 cell stage fertilized egg of Tg(HUC:GAL4;UAS:synaptophysin-GFP) lead to mosaic UAS-mCherry caax incorporation in some cells, while in the transgenic fish line Tg(HUC:GAL4;UAS:synaptophysin-GFP) synaptophysin fused GFP is expressed at the presynaptic site in presumably all HUC expressing cells (arrowhead direction corresponds to transcription and translation processes). As a result, mCherry mosaic expression will be visible in cell membranes in occasional neurons. (2) During selection process only zebrafish with labelled neurons in the caudal fin were considered for further analysis. Selection was performed at 5 dpf to ensure developed branching of caudal fin neurons. Zebrafish with mCherry fluorescence in the caudal fin were manually collected and used for the analysis. (C) *In vivo* confocal imaging of 5 dfp mCherry-caax-XHUC:GAL4:synaptophysin-GFP zebrafish caudal fin. Magnified inset shows lack of punctiform artifacts. This feature makes our mCherry reporter line preferable for neuronal branching reconstruction and thus was used in all experiments to visualize generation of neuronal trees.

#### S3.4 Imaging of live animals

The expression of mCherry in cell membranes of zebrafish neurons is not uniform, including in the region of the caudal fin, therefore we first screened multiple animals and selected fish which presented mCherry positive signals in the caudal fin. 5dpf fish were anesthetized with tricaine (MS-222, Sigma) in final concentration 200 ug/nil in E3 PTU treated medium. Then 5 fish samples per dish were immobilized in 500 ul of 0.5%. low melting agarose (LMA, Sigma), supplemented with tricaine (200 ug/ml) and placed laterally on glass bottom microwell dish (MatTek, uncoated, 35mm) using tungsten forceps. After complete polymerization of LMA (40-60 min at room temperature), the droplets containing live fish were covered with Tricaine supplemented with E3-PTU medium to prevent desiccation of the immobilized fish during imaging. Confocal images were acquired using Z-stacks with a Zeiss LSM 800 confocal microscope equipped with Diod lasers 405 inn, 488 inn, 555 inn and 639 inn, Plan Apochromat lOx, Plan Apochromat 20x, and C-Apochromat 40x. Images were processed in Bitplane Imaris 8.0 and exported as .tiff files for further analysis.

#### S3.5 Reconstruction of neuronal filaments

##### S3.5.1 Initial (manual) reconstruction of the filaments

Arborization trees of all visible neurons-the transient Rohon-Beard sensory neurons located in the dorsal spinal cord and innervating the caudal fin integuments [15] – were reconstructed using the pipeline described below. Raw images acquired on live animals were exported from ZEN software to Bitplane Imaris 8.0. For initial reconstruction, Bitplane Imaris tool “Filaments” was used. Resulting images are referred to as “filament trees” in the following. For each image multiple filament trees were acquired. Each tree corresponds to a unique neuron arborization. Each filament tree was then manually analyzed, using native mCherry fluorescence channel as a reference, to eliminate false connections between branches. After manual correction, the tools “smooth filaments” and “center filaments” were applied to further co-localize obtained reconstructions with fluorescence signal. Filaments with confirmed branching pattern and no cross-connectivity artifacts were taken to the next step of analysis (see below for details). All filaments which could not be clearly traced were eliminated from the further analysis. 2D Images of the selected filament trees were exported as separate .tiff files and transferred to Imaged software. The same set of tools was applied to all filaments: conversion to 8-bit black and white image, skeletonize, analyze skeleton.

###### Limitations of the experiment

Prior to the manual reconstruction of the filaments, we performed raw image quality assessment, based on the following parameters. Neurons chosen for reconstruction had a minimal optical overlapping with its neighbors. All raw images were acquired using the same confocal microscopy settings to ensure uniform data resolution. We note that, however, based on the pinhole settings, we could not reliably distinguish between two dots if the Z distance was less than one pinhole, which was set for 1 AU (airy unit) and equal to 1.75 um at 20x objective and to 6.45 at lOx objective. Therefore, any neurites (from two different neuron trees) closer than this distance were discarded from the analysis, creating a small loss in the reconstruction. In a sufficiently dense region of the fin, this loss was evaluated post-hoc to amount to 8.9% of the overall length of all mCherry positive axon arbors (using Imaris tool “filament statistics”). Overall, considering these limitations, a total number of 8 filaments from about 50 fish scanned were qualified for analysis.

##### S3.5.2 Coarse-grained reconstruction of the filaments

To obtain quantitative information on the branch numbers, branch lengths and probabilities of branching events, we needed to convert the manually reconstructed filament images into skeletonized datasets to extract coordinates. To achieve this, we first conducted the following analysis using custom scripts in Python: The manual reconstruction images of filament and fin borders were loaded and separated according to the channel information. After binarization, images were skeletonized with Lee’s algorithm [16] implemented in scikit-image (v. 0.17.1) [17]. The Skan module (v. 0.9) [18] yielded a vector representation of filaments and border outlines. Finally, vector size and position were adjusted to correct for differences in input image resolution.

**Figure S8:**
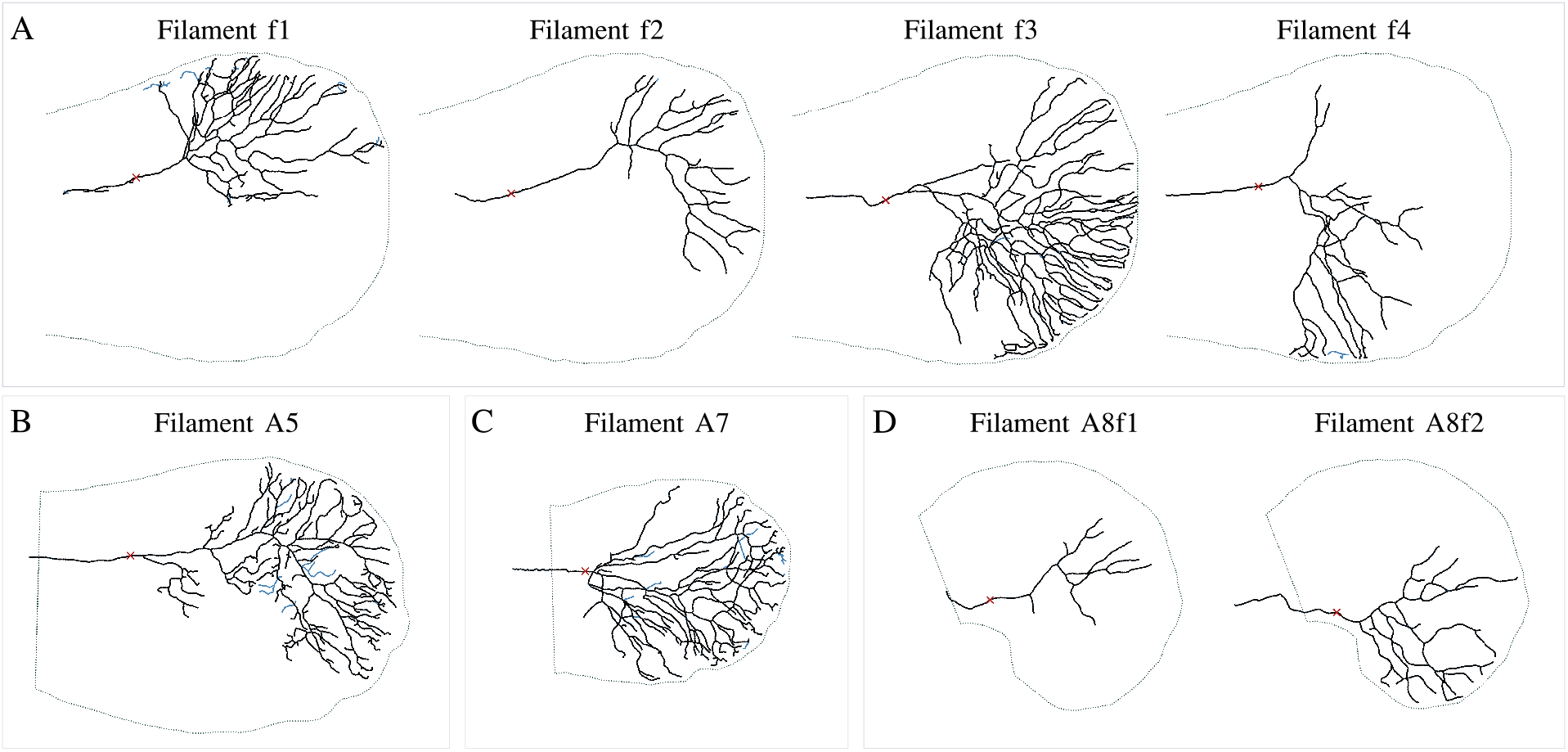
Reconstruction of experimental data after applying the coarse-graining algorithm. The data points shown in black correspond to the final coarse-grained coordinates that are used in the analysis, whereas the skeletonized data from the reconstructed images are shown in blue. Filaments from the same fish are displayed in the same box. The fin edges were identified manually and are indicated by the thin dashed lines. Data acquisition corresponds to 5 dpf for all 8 samples.

After the skeletonization with the Skan module, we still had to define hierarchical trees with a well-defined orientation for the branches. Moreover, the skeletonized networks now consisted of elementary vectors that only attained a discrete set of local angle values (integer multiples of *π*/4) and had a length of about a single pixel. We therefore developed an algorithm in order to coarse-grain the networks starting from an initial point of origin. The resulting networks then consisted of discrete vectors of a pre-defined stepsize *ℓ*, which had local angles with a finer distribution of values.

For the coarse-graining algorithm, we first label all branching points in the network by identifying the three-valent vertices. We then start the coarse-graining loop by defining an “active tip” which is the vector closest to the origin. This active tip will evolve by “scanning” an area of a certain radius *R* for the underlying skeletonized data points and taking discrete steps (of average size 〈*s*〉 determined mainly by the scanned radius *R*) according to the following rules:

i. *Transition towards branching point*: If there is a branching point in the scanned area of radius *R*, the active tip will jump towards this point in the next time step.
ii. *Elongation*: If there is no branching point in the scanned neighborhood, the active tip will try to move forward with respect to its current polarity by defining a “polarity cone” that provides a radial slice of the scanned neighborhood in the direction of the tip, and selecting the furthest data point of the underlying skeletonized network within the polarity cone in the next time step.
iii. *Branching*: If the active tip itself is a branching point (which will necessarily be the case for a tip that undergoes the transition (i) above), it will search for two data points as progenies and produce two active tips at these coordinates in the next time step. The triangle connecting the two progenies and the active tip is required to have a minimal angle value at the vertex of the active tip in order to prevent branching events into the same branch of the underlying network.
iv. *Annihilation*: If the above conditions are not fulfilled, the active tip will become inactive in the next time step, *i.e*., it will not be iterated further in the loop.

**Figure S9:**
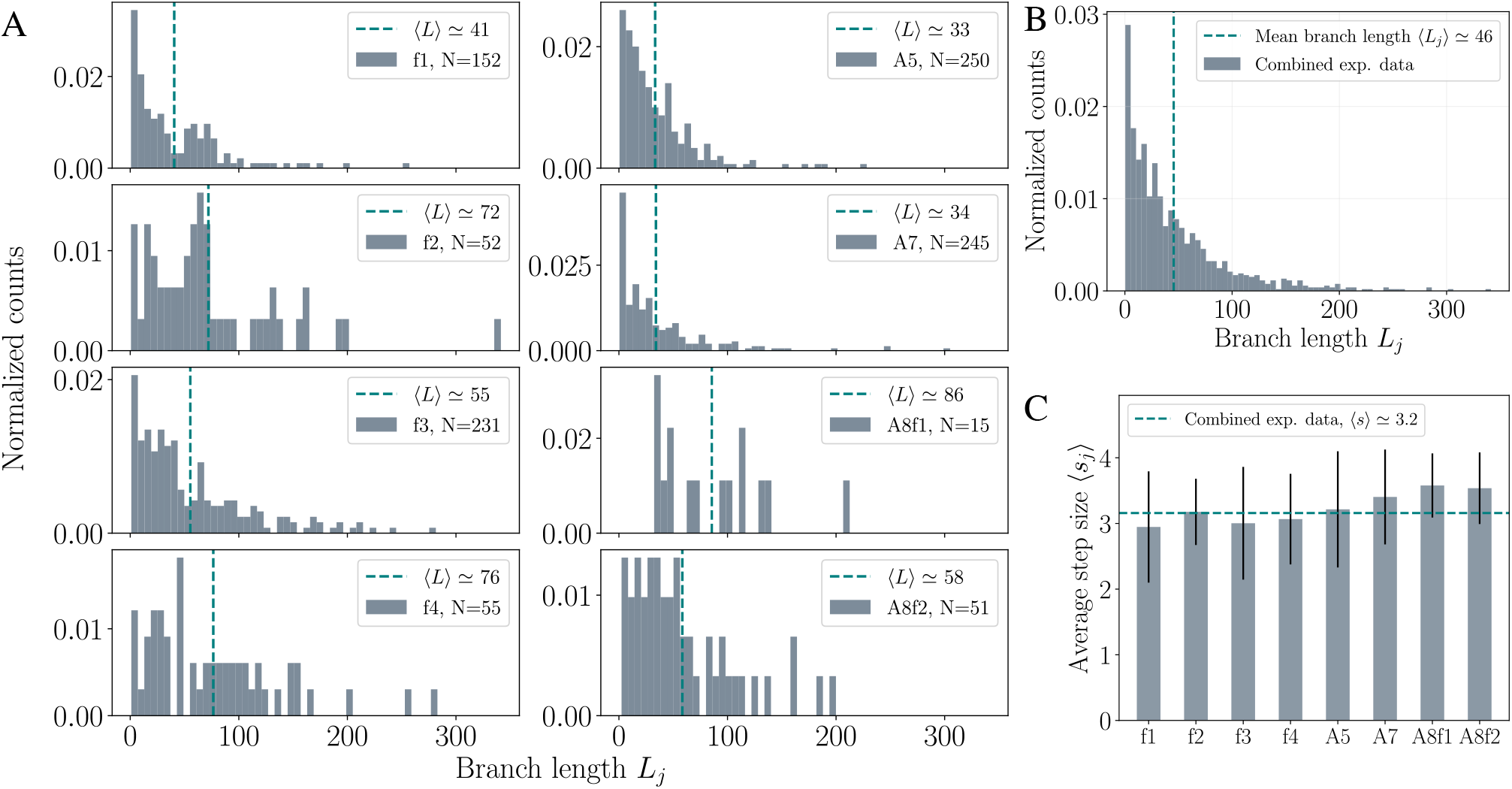
(A) Normalized histograms of branch lengths (in pixels) of individual filaments after the reconstruction of experimental data with the coarse-graining algorithm. The number of branches *N* for each sample are indicated in the inset. Dashed vertical lines represent the mean branch length 〈*L_j_*〉 with *j* = 1,…, *N* for the individual samples. (B) Normalized histogram of branch lengths for the combined data from all *n* = 8 filaments. Dashed vertical line represents the average branch length 〈*L_j_*〉 of the combined data. (C) Bar chart visualizing the individual average step sizes of the coarse-graining algorithm for each sample. The mean branch size 〈*L_j_*〉 (dashed line’ in B) and the average step size 〈*s*〉 of the combined dataset (dashed horizontal line) can be used to estimate the average number of steps in a typical branch via 〈*L_j_*〉/〈*s*〉 ≃ 14.

In some cases, two active tips start “invading” the same branch due to noisy regions in the original raw images. In order to prevent such events, we implement the additional rule that when an active tip “moves” on an already processed data point, it will immediately terminate. In the final data processing step, we then revise the data of the corresponding branch such that it acquires the generation label and orientation of the “older” active tip that has a longer ancestral lineage.

This set of rules closely resembles the simulation setup for BARWs and provided a hierarchical tree for each sample from the experiments. After the coarse-graining loop, each processed datapoint was assigned a generation label and a local angle value determined by the vector connecting its positions at time *t* and in the previous time step *t* – *τ*. The resulting coarse-grained coordinates for the *n* = 8 networks analyzed here are shown in Fig. S8 in black, where the underlying skeletonized networks are presented in blue.

**Figure S10:**
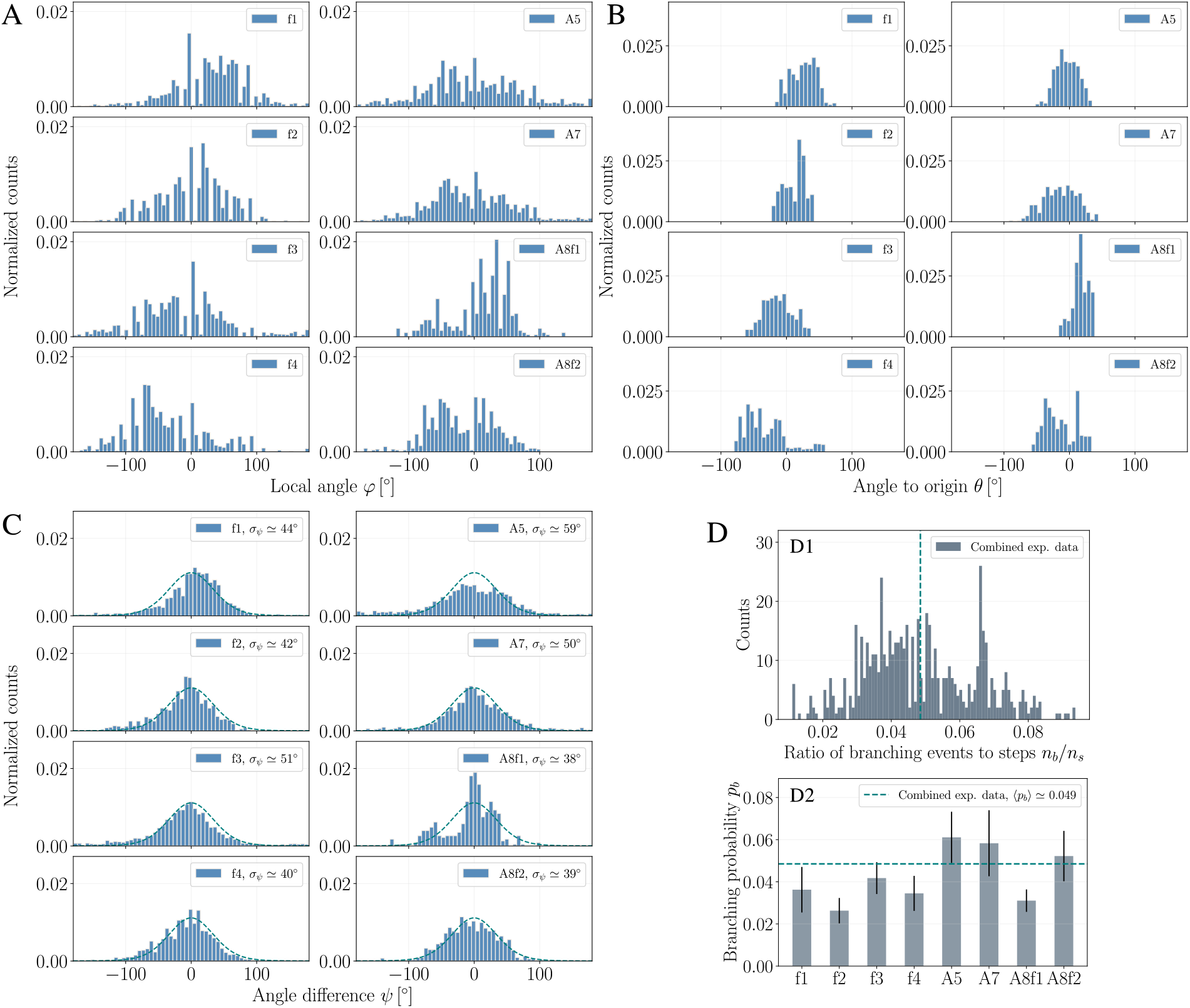
(A-C) Angle distributions obtained from experimental data. Normalized histograms of the (A) local angle *φ*, (B) angle to origin *θ*, and (C) angle difference *ψ* values for the individual filaments. Despite significant fluctuations, the latter histograms can be well-described by the von Mises distributions (dashed lines) with an estimated branching probability of *p_b_* = 0.05 and a field strength of *f_c_* = 0.6. (D) Estimation of the branching probability *p_b_* for the experiments data from the ratio of the number of branching events *n_b_* to the number of steps *n_s_* until the endpoint (“leaf”) of each branch lineage is reached. (D1) Histogram of the ratio *n_b_/n_s_* for the combined data from *n* = 8 filaments. (D2) Estimates for *p_b_* for the individual filaments. Average branching probability of the combined data is estimated as 〈*p_b_*〉 – 0.049 (dashed horizontal line). Error bars indicate the standard deviations of the individual distributions.

#### S3.6 Analysis of experimental data

##### S3.6.1 Distribution of branch lengths

A measure that is intimately linked to the probability of branching is the average branch length 〈*L_j_*〉 of a filament with *j* = 1,…, *N*, where *N* denotes the total number of branches in the tree. For a network with a high *a priori* branching probability, one would expect to find short branches on average due to the frequent branching events. To quantify this, we estimated the branch lengths *L_j_* by calculating the total length Σ*_k_s_k_* of all branch segments (discrete steps of size *s_k_*) between the starting and end points of each branch. The normalized histograms of branch lengths for the individual filaments are displayed in Fig. S9(A). Importantly, the combined data from *n* = 8 samples showed an exponentially decaying tail (see Fig. 2(F) in the main text) with an average branch length of 〈*L_j_*〉 ≃ 46, see Fig. S9(B). This is what is expected in a stochastic branching process, validating a key assumption of the framework of BARWs. We also calculated the average step sizes corresponding to single steps of the coarse-graining algorithm described above, and obtained a mean step size of 〈*s*〉 ≃ 3.2 from the combined dataset, see Fig. S9(C). The mean step size can be used to determine the normalized branch lengths 〈*L_j_*〉/〈*s*〉 ≃ 14 to compare the lengths with the simulation data (as shown in Fig. 2F in the main text).

**Figure S11:**
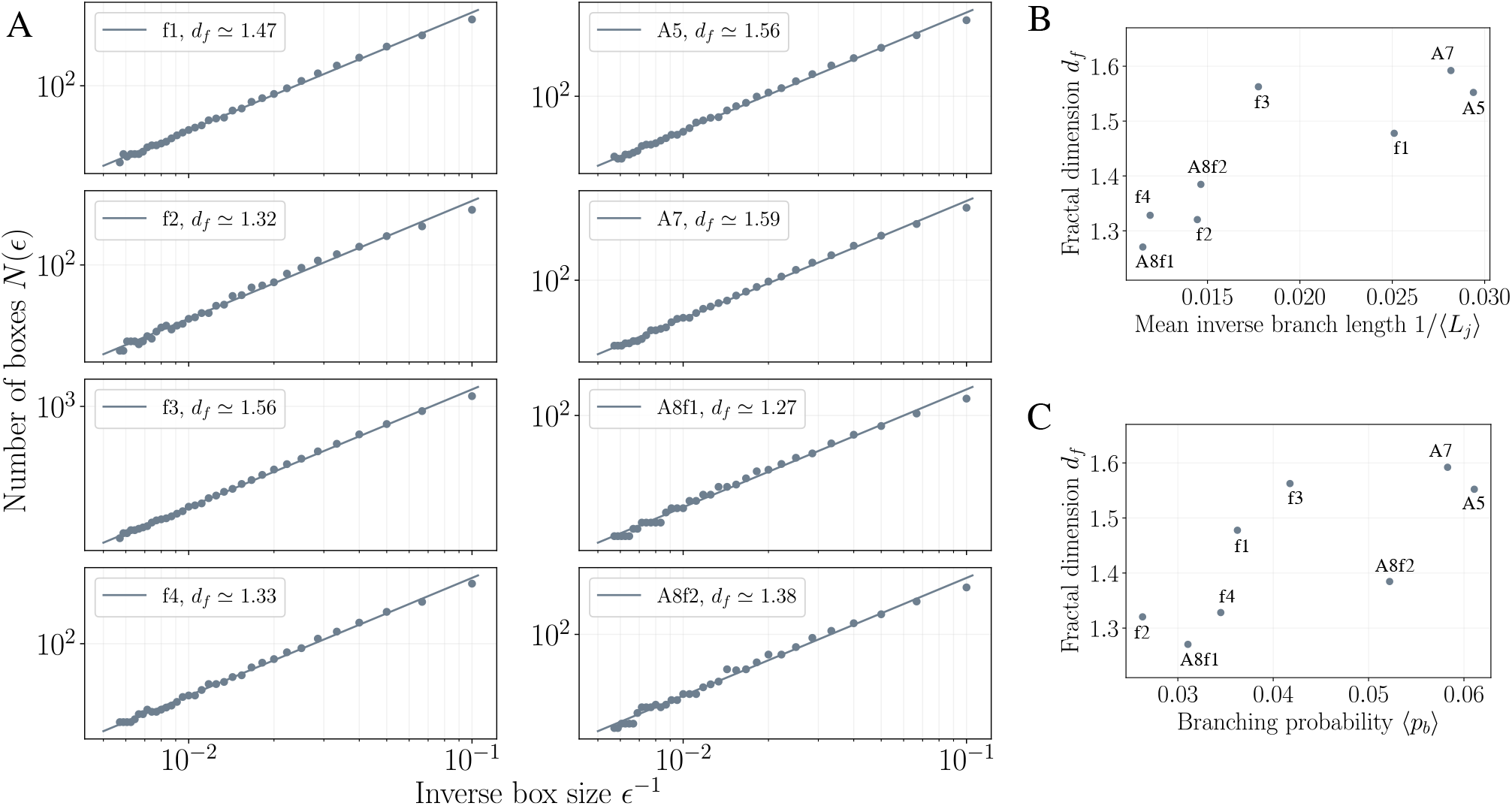
(A) Fractal dimensions of the individual filaments obtained from experimental data using the box-counting method: Box sizes of varying sizes (from *ϵ*_min_ ≃ 3〈*s*〉 to *ϵ*_max_ ≃ 3〈*L_j_*〉) are used to count the number of nonempty boxes *N*(*ϵ*) for individual networks. (B) Estimated fractal dimensions correlate with the mean inverse branch lengths 1/〈*L_j_*〉 and (C) with the estimated branching probabilities 〈*p_b_*〉 of the individual samples. Note that the latter values arc not strictly proportional to the inverse branch lengths.

##### S3.6.2 Estimation of the branching probability

A key parameter to determine for the comparison of our theoretical results with the experimental data is the branching probability *p_b_* of the networks. One can estimate *p_b_* from the measured distributions of branch lengths because the average branch length depends in general inversely on the branching probability, *i.e*. 〈*L_j_*〉 ∝ 1/*p_b_*. However, due to the frequent annihilation events the distributions of branch lengths in fact underestimate the average branch length, and thus lead to a high branching probability *p_b_* that generate networks qualitatively different from the experimental observations. We therefore decided to estimate *p_b_* by directly counting the number of branching events *n_b_* for each lineage and taking its ratio to the total number of branch segments (steps) *n_s_* for that lineage. The branching probability *p_b_* of a network then corresponds to the average ratio 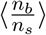 from all lineages of branches ending at a leaf. Fig. S10(D) displays the normalized histograms for the ratio 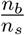 obtained from the combined dataset of *n* = 8 filaments (DI) and the estimates of *p_b_* for the individual samples (D2). The average branching probability obtained from the combined data of 〈*p_b_*〉 ≃ 0.05 was finally used to generate the branching networks of the BARW simulations and to compare the results on angular alignment by determining the diffusion and mobility coefficients given in Eq. (S18).

##### S3.6.3 Angle distributions and fractal dimensions for individual samples

To obtain the angle distributions we first needed to clarify the coordinate of the origin for each individual sample. Because the initial branch of some filaments was located in the spinal cord outside of the fin region, we did not take the starting point of the initial branches as the origin. Instead, an alternative choice which had the advantage of being systematic was to first locate the boundary of the fin tissue with respect to the anterior-posterior axis, and then fix a central point that has the same distance to the boundary for all samples. After this identification, angles to origin *θ* and angle differences *ψ* could be calculated for each node in the coarse-grained networks. The normalized histograms for the local angles *φ*, angles io origin *θ*, and the angle differences *ψ* for the individual filaments are shown in Fig. S10(A-C). The histograms for the local angle *φ* and the angle to the origin *θ* attained rather irregular shapes, whereas the histograms for the angle differences *ψ* for most of the individual samples could be well-approximated by the analytical predictions (Fig. S10(C), dashed lines). Importantly, the prediction held for both denser and sparser networks, which is a key prediction of the model, and wouldn’t hold if for instance the directionality was emerging from the repulsion between a dense network of tip/branches (see Fig. S3(D)).

Finally, in Fig. S11 we display the fractal dimensions for the individual samples obtained by the box-counting method. For this purpose, we selected box sizes gradually increasing from *ϵ*_min_ ≃ 3〈*s*〉 to *ϵ*_max_ ≃ 3〈*L_j_*〉 to describe a well-defined scaling behavior (*i.e*. to omit boundary effects arising from too large or too small boxes). The fractal dimensions showed a rather robust correlation with the inverse of the mean branch lengths and with the branching probability, even though these two measures were not strictly proportional for all samples, see Fig. S11(B) and (C).

